# Parallelization of particle-based reaction-diffusion simulations using MPI

**DOI:** 10.1101/2024.12.06.627287

**Authors:** Sikao Guo, Nenad Korolija, Kent Milfeld, Adip Jhaveri, Mankun Sang, Yue Moon Ying, Margaret E Johnson

## Abstract

Particle-based reaction-diffusion models offer a high-resolution alternative to the continuum reaction-diffusion approach, capturing the discrete and volume-excluding nature of molecules undergoing stochastic dynamics. These methods are thus uniquely capable of simulating explicit self-assembly of particles into higher-order structures like filaments, spherical cages, or heterogeneous macromolecular complexes, which are ubiquitous across living systems and in materials design. The disadvantage of these high-resolution methods is their increased computational cost. Here we present a parallel implementation of the particle-based NERDSS software using the Message Passing Interface (MPI) and spatial domain decomposition, achieving close to linear scaling for up to 96 processors in the largest simulation systems. The scalability of parallel NERDSS is evaluated for bimolecular reactions in 3D and 2D, for self-assembly of trimeric and hexameric complexes, and for protein lattice assembly from 3D to 2D, with all parallel test cases producing accurate solutions. We demonstrate how parallel efficiency depends on the system size, the reaction network, and the limiting timescales of the system, showing optimal scaling only for smaller assemblies with slower timescales. The formation of very large assemblies represents a challenge in evaluating reaction updates across processors, and here we restrict assembly sizes to below the spatial decomposition size. We provide the parallel NERDSS code open source, with detailed documentation for developers and extension to other particle-based reaction-diffusion software.

## 1. INTRODUCTION

Reaction-diffusion (RD) models are widespread tools for quantifying the dynamics of nonequilibrium systems in physics^1^, chemistry^2^, engineering ^3^, and quantitative biology^4,5^. By capturing interactions between species via rate-parameterized reactions as the species diffuse in 3D, 2D, or 1D environments, they predict the emergence and time-evolution of spatial patterns and order. From early studies on morphogenesis by Alan Turing in the 1950s^6^, the reactions introduce nonlinearities into the differential equations and thus exact solutions must be determined numerically. Continuum RD models can be solved relatively efficiently on computers via discretization of space and time using finite element methods, with commercial and academic options like COMSOL® or Virtual Cell^5^. However, continuum RD models lack particle resolution or stochasticity, limiting their ability to capture the integer-valued copy numbers of physical objects or the nanoscale self-assembly processes that often represent fundamental steps in pattern formation. While lattice-based RD models (reaction-diffusion master equation) do capture integer-valued copy numbers and stochasticity^7–12^, they are well-mixed within lattice voxels, thus they do not capture the volume or structure of individual species, preventing simulations of nanoscale self-assembly. Self-assembly and self-organization are essential elements of chemical systems and living systems, governing function across multiple scales. Particle-based reaction-diffusion models thus provide an important higher-resolution method for studying the dynamics of nonequilibrium systems as they depend on the stochastic interactions and higher-order assembly of individual reactive species. The increased resolution of particle-based RD comes at a significant computational cost^4^, however, and current parallelization strategies to accelerate timesteps are limited to single-node GPU-based methods^13,14^, which therefore cannot take advantage of less expensive and ubiquitous CPU resources. Accelerating particle-based RD methods not only improves the efficiency of simulating processes over long timescales, but by shortening simulation times significantly, allows one to take advantage of parameter optimization techniques like automatic differentiation that can learn model parameters^15^ but are limited by the cost of simulation times. Here we present a parallel implementation of the particle and rigid-body structure-resolved RD software NERDSS^16^ using the message passing interface (MPI)^17,18^, supporting linear scaling over several computer nodes for large systems.

The NERDSS software is uniquely designed to handle ‘molecules’ defined as rigid collections of spherical sites that can self-assemble into higher-ordered structures^16,19^, building off single-particle RD algorithms for dynamics in 3D^20^, 2D^21^, and transitions between^22^. The software thus shares the same basic structure and order of operations of other particle-based RD software, allowing most of the parallelization methods used here to be directly transferred to other tools like Smoldyn^23^, MCell^24^, ReaDDy^25^, GFRD^26^, and SpringSaLaD^27^. All particles are updated over fixed timestep intervals, with positional displacements sampled according to Brownian motion. A time-consuming calculation in each timestep is the evaluation of binding reactions between particle pairs, dependent on their probability of collision or co-localization in a small volume. While the exact details of these reaction probability calculations vary between algorithms^4^, they are relatively short-ranged because collision probabilities fall to zero with increasing separation, dependent on the timestep and the particle diffusion constants^20^. Thus the spatial or domain-decomposition across processors that we implement here takes the best advantage of the locality of the most expensive binding computations. NERDSS encodes models using commonly accepted rule-based standards for reactions^28^, including formatting for spatial models, supporting portability of data structures. A distinct challenge for NERDSS and other multi-site reaction-diffusion methods like SpringSaLaD is that the formation of larger assemblies, whether disordered (SpringSaLaD)^27,29^ or for NERDSS, structured assemblies like spherical cages^30,31^, lattices^32^, surface clusters^33^, and filaments^34^, does create longer-range coupling within single assemblies. For large rigid bodies, they move as a unit, requiring a single processor to control displacement despite potentially spanning multiple spatial domains. We address the need for displacement control with a single processor here via added communication steps, which currently places a limit on the maximal assembly size.

Our parallel implementation of particle-based RD takes advantage of successful techniques used for decades in the molecular dynamics (MD) community^35–43^, as MD simulations similarly propagate particle (atom) positions in 3D space via differential equations. MD simulations also evaluate pairwise interactions between particles within a cutoff distance, and MPI implementations for MD software^37–39^ therefore use a spatial or domain decomposition of the system, where contiguous regions of physical space are assigned to separate processors to partition the particles. We similarly implemented a spatial decomposition including shared or ‘ghosted’ regions, where copies of particles owned locally by one processor are duplicated onto neighboring processors to facilitate computation of particle-particle interactions across the domain boundary, minimizing repeated communication steps and thus overall communication delay^44–46^. A primary difference between MD and RD simulations is that all particles interact with one another in MD simulations and at higher densities, whereas in RD, only reactive partners typically evaluate pairwise interactions. MD thus typically has far more pairwise interactions to evaluate and at a higher computational cost per evaluation, making the total computational cost per timestep higher; this supports better scaling behavior up to thousands of processors. MD is less sensitive to variations in parameterizations, as the particle densities and number of computational evaluations does not change from one atomistic (or coarse-grained) system to the next. RD simulations can support a much wider range of particle densities and reaction networks, meaning the cost and the scaling efficiency of RD is problem dependent. Nonetheless, we show here that even lower density systems can benefit from parallelization.

While there are several options to accelerate scientific software across CPUs or GPUs, we focused here on an MPI distributed memory strategy across multiple CPU nodes because it is an efficient, scalable, and economical approach. Particle-based simulators for MD historically all adopted MPI to exploit the thousands of nodes available on supercomputers^35,37,38^. While modern MD packages typically also provide hybrid modes augmented to also use GPUs, they must then compete for these scarcer GPU-based resources, and although they show excellent performance on a single node, they do not scale as well across multiple nodes^39,47,48^. These trade-offs mean that both MPI-based and GPU-based MD software are actively used for large-scale simulations. These same trade-offs exist for particle-based RD, but with the additional factor that RD is less optimal for GPU parallelization compared to MD, because its operations and data-access patterns can vary over the course of a simulation due to creation and destruction of particles and highly heterogeneous particle distributions undergoing potentially new reactions per step. Nonetheless, two particle-based RD software packages have used custom GPU implementation^49^ or leveraged GPU-accelerated MD frameworks for RD operations to successful accelerate performance on a single GPU node^50^. Our MPI implementation of the particle-based NERDSS software is thus the first such CPU-parallelized version that we know of, supporting scaling across multiple CPU nodes and a general strategy applicable to other particle-based RD tools.

In this paper, we detail the implementation and algorithm modifications required to parallelize the C++ NERDSS software^16^ using MPI, shown schematically in Fig. 1. We address the challenges of efficiently distributing molecules and complexes across the processors, managing interactions and reactions that span processor boundaries, and ensuring data consistency and accuracy in the presence of complex, multi-molecule assemblies. To minimize additional development, we retain the serial NERDSS computing kernel as much as possible, documenting necessary modifications for data communication. We validate the accuracy of parallel NERDSS by simulating simple bimolecular reactions in various spatial configurations (3D, 2D, and 3D to 2D) and comparing with theory and serial NERDSS. We then extend our validation to more complex macromolecular assemblies and hexagonal lattice self-assembly. Comprehensive performance benchmarks for strong scaling and weak scaling are conducted for all simulated systems to quantify the scalability and efficiency gains achieved through parallelization. We conclude by discussing the current limitations of our parallel implementation and outlining future development directions to further enhance its capabilities and performance. NERDSS and parallel NERDSS are both accessible open-source under the GNU General Public License at github.com/mjohn218/NERDSS.

**Fig 1.**
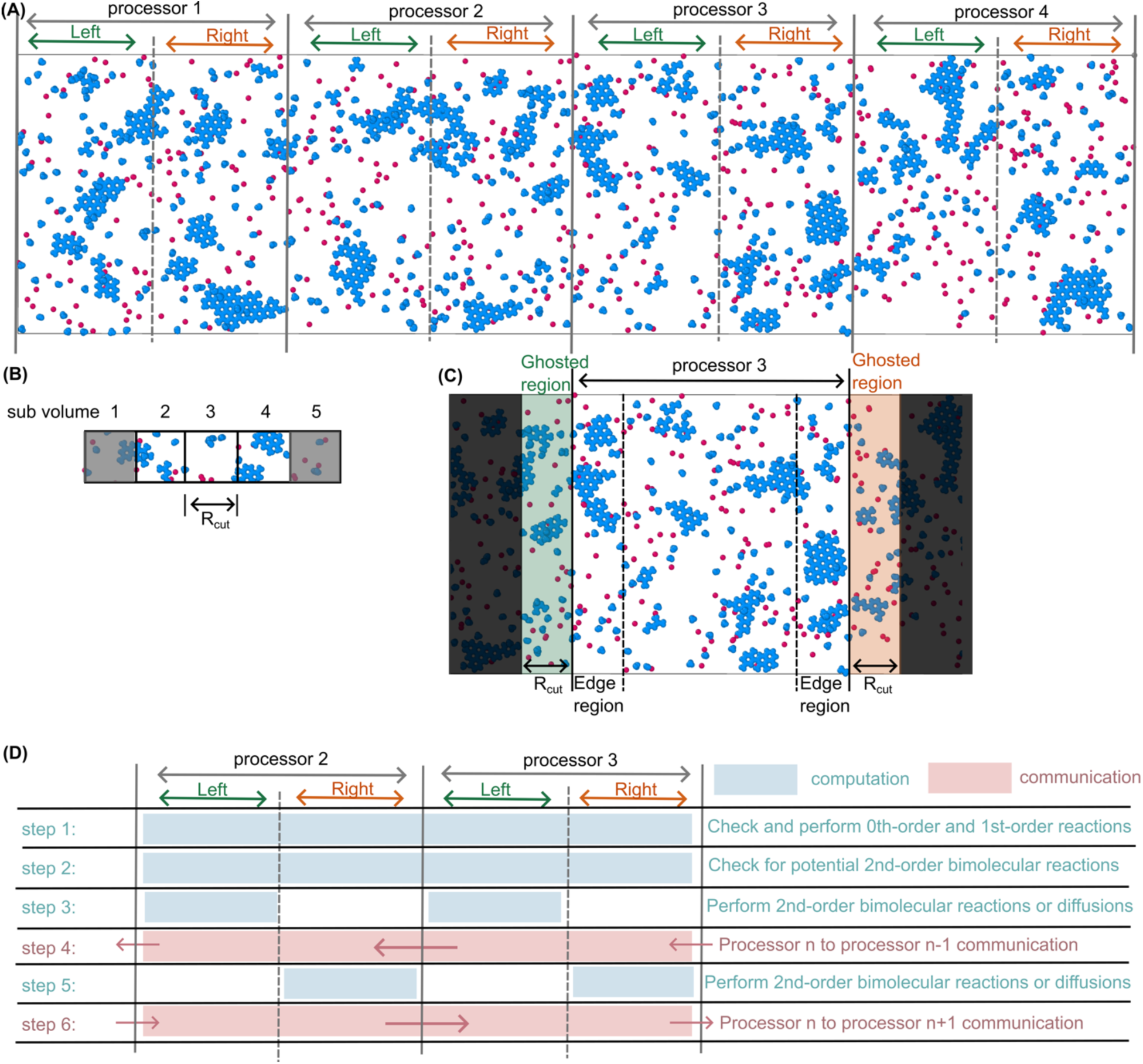
Overview of the parallel NERDSS implementation. (A) The 3D simulation space is divided only along the x-axis to distribute the workload across multiple processors. Each processor’s region is divided into left and right halves to synchronize communication with concurrent computation. (B) Within each processor, sub-volumes are defined to optimize the computation of pairwise binding probabilities, with each sub-volume’s length in x, y, and z defined to be at least the length *R*_cut_(Eq. (3)) to ensure all potential binding events are evaluated within Δ*t*. We illustrate sub-volumes here along x, where molecules in sub-volume 3, for instance, can only interact with molecules in the same sub-volume 3 or in the neighboring sub-volumes 2 and 4. (C) Ghosted regions are used to manage interactions across the processor boundaries by duplicating molecules from the neighboring processors (2 and 4) to be evaluated for pairwise interactions within processor 3. The edge regions are the sub-volumes within the processor that are at its boundary, adjacent to the ghosted regions. The edge regions on one processor are thus ghosted regions on another processor. (D) A staggered computation in each half is used to reduce idle time and ensure all processors work simultaneously between communicating with left and right neighbors. Processor *n* perform 0^th^ and 1^st^ order reactions (step 1), calculates 2^nd^ order reaction probabilities (step 2), executes 2^nd^ order reactions and diffusion events in the left half (step 3), communicates border molecules information to its left processor *n*-1 (step 4), computes its right half (step 5), and communicates with its right processor *n*+1 (step 6).

## 2. METHODS

### 2.1. Background on the NERDSS particle-based reaction-diffusion software

NERDSS simulates stochastic reaction-diffusion dynamics with interactions between spherical particles and/or rigid bodies defined by multiple spherical interaction sites. The simulations are bounded within either a sphere or a rectangular box, with reflecting boundary conditions by default. Membranes are modeled in two ways: either as a reflecting surface that houses diffusing particles (e.g., lipids) restricted to 2D, or as a partially adsorbing surface, implemented as an implicit lipid model where lipid densities change to reflect binding events^22^. The diffusional propagation of particles and the evaluation of reaction probabilities is performed via the Free-propagator reweighting algorithm (FPR)^20^, described further below. The algorithm retains exact association rates for pairs despite using a free-propagator position update by applying trajectory reweighting to reaction probabilities^20^. The algorithm applies equally to reactions on a 2D surface^21^, and between 3D and 2D surfaces^22^.

To briefly describe the reaction-diffusion propagation, each particle or rigid body ‘molecule’ in the many-body system undergoes either a diffusion or a reaction in each fixed timestep Δ*t*. Reactions are evaluated first. For first-order reactions with rate *k* (such as dissociation events), the number of species *N* capable of performing that reaction is tracked, and the number of events *n* to perform in the timestep is computed via the probability *k*Δ*t* by sampling from a binomial distribution, which is faster and equally accurate to computing reactions for each specie one-at-a-time. The most time-consuming calculations involve updates between particles that are close enough (defined below) to collide and thus react with one another via 2^nd^ order reactions. Maximally one interaction site on a rigid-body molecule can undergo reaction in a timestep. For a larger complex that consists of multiple assembled molecules, distinct molecules can each undergo a reaction. For all particles and molecules that do not perform reactions, they are propagated using Brownian updates based on their translational and rotational diffusion. Diffusion constants scale with the size of the complex based on the Einstein-Stokes relations, as each rigid-body or complex moves as a single unit relative to its center of mass. New positions of all interacting sites must avoid overlap (exclude volume) with all reactive partners. When binding reactions occur between multi-site molecules, the orientations they adopt is defined by user-specific angles to control the structure of the multi-protein assembly (e.g. filament, spherical lattice). If association between sites on two molecules produces steric overlap of other sites within the complex, the binding event is rejected to ensure volume is excluded per molecule and physically realistic structures are generated. A pseudo-code is provided in the SI.

### 2.2. Pairwise reactions and domain size limits

For parallelization of NERDSS, the key steps that require communication between nodes are the evaluation of pairwise reactions across the processor boundaries, and the placement of particles and rigid bodies to avoid overlap following Brownian updates. For many-body systems, the FPR algorithm treats each pair independently. To support large time-steps while still resolving collisions between a pair of particles 1 and 2, FPR uses the Green’s function (GF) solution, *p*(*r*, Δ*t*|*r*_0_) for finding the diffusing pair at separation *r* after Δ*t* given an initial separation *r*_0_. The pair are subject to a radiation boundary condition at the collision radius *σ*, parameterized by the intrinsic rate *k_a_*. The spatial integral over this GF provides the probability of a reaction occurring in Δ*t*, which in 3D is given by:

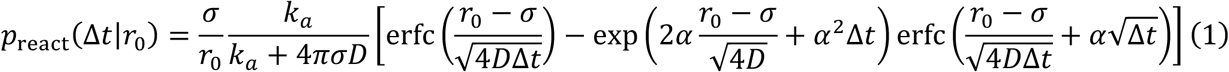

where *D* = *D*_1_ + *D*_2_and *⍺* = D√*D*/*σ*F(1 + *k_a_*/(4*πσD*)). The reaction probability thus depends on the diffusion constants of the species, and if the reaction is occurring between sites on multi-site molecules, we approximate the impact of rotational diffusion into an effective diffusion constant that retains accurate reaction probabilities^19^. Forces can be introduced into this RD framework, as shown previously^20^. This is not a standard element of NERDSS, however, because it changes the reaction probabilities from analytical formulas like Eq. 1, rendering them numerically expensive to compute.

Reaction probabilities drop to zero with increasing separation *r*_0_ (Eq. 1), thus setting the maximal cutoff where pairwise reactions must be evaluated. Instead of inverting Eq. 1 to solve for the separation where *p*_react_(Δ*t*|*r*_0_) drops to zero, we use a simpler definition of a cutoff based on the average displacement due to diffusion, scaled by 3. This approximation works because the particles cannot react (*p*_react_(Δ*t*|*r*_0_) → 0) if they cannot diffuse to contact, which is controlled by Δ*t*, *D*, and σ. This cutoff is defined for each bimolecular reaction *m* via:

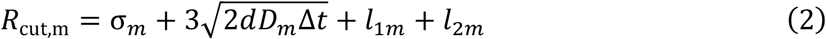

where *d* is the dimensionality of space and *l_im_* (*i* = 1 *or* 2) accounts for any displacement of the reactive site from the molecule center-of-mass. The maximal cutoff distance across the entire system is then given by:

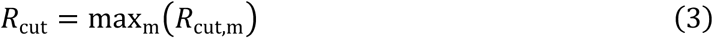

This *R*_cut_ is important; it establishes a limit on the minimal domain size that needs to be shared with neighboring processors to ensure that all reaction pairs are evaluated.

We note that the maximal timestep Δ*t* is also restricted by either the fastest timescale in the reactions or by the density of the system. The latter consideration is typically much more restrictive and arises because the FPR algorithm, to retain exact association probabilities, assumes a particle can only react with at most one partner in Δ*t*. Based on the density of reactive partners and *R*_cut,m_, the timestep for a given reactant 1 with *N*_1_ copies due to binding to partner 2 with *N*_2_ copies in a box with side length *L* is maximally:

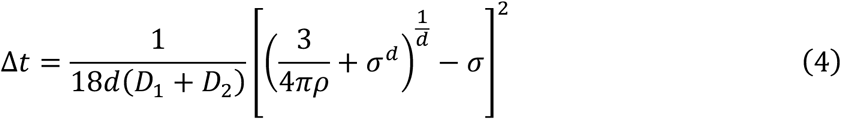

where 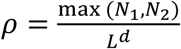. The limiting timestep for the entire system is then the shortest for all reaction pairs. Due to density fluctuations, a particle can still encounter more than one partner in its reaction cutoff. Each reaction is evaluated independently. Our extensive results on many-body dynamics compared to theory^4^ indicate that deviations from the pairwise approximation have negligible impact on the accuracy of the dynamics; even when using Δ*t* values 10x larger than this conservative estimate, the results are still accurate.

### 2.3. Sub-volume partitioning in serial NERDSS

Even within serial NERDSS or on a single processor, we take advantage of the short-range nature of the interactions by partitioning the full region into rectangular sub-volumes that have a minimal side length of *R*_cut_in each dimension x, y, and z (Fig 1). Instead of looping over all particle pairs, we thus only evaluate particle pairs within your own sub-volume and 13 neighboring sub-volumes^16^ (Fig S1). There is an optimal number of sub-volumes, as having too many can exceed the number of particles within the simulation volume. We therefore place an upper limit on total sub-volumes to enhance efficiency, while using the minimal side length of *R*_cut_in the x dimension (Fig S1).

### 2.4. Parallelization strategy via spatial decomposition

We implemented a spatial decomposition approach in parallel NERDSS that divides the full simulation space along the x-axis into regions each assigned to a different processor (Fig 1A). This simplifies the communication protocol significantly compared to a full parallelization in x, y, and z, which would be the focus of future work. We chose the x-dimension because the membrane can create a significant density imbalance (and thus load imbalance) in the z-dimension when molecules partition from 3D to the surface (at z=0).

NERDSS stores coordinates of all particles or molecules in the ‘Molecule’ data structure. This structure also tracks the states of all binding sites within the molecule (e.g., free or bound). NERDSS maintains a separate ‘Complex’ data structure that changes size depending on the number of distinct assemblies and/or particles in the system. The Complex structure tracks the indices of all the molecules that make up the complex, along with the center of mass coordinates of the entire complex. Each Molecule and Complex structure are assigned to a single processor based on their spatial coordinates. However, all Molecules in sub-volumes at the edge of each processor’s region are sent and duplicated to the neighboring processor as their ghosted regions (Fig 1C). These duplicated or ‘ghosted’ molecules are essential for evaluating bimolecular reactions between processors. Likewise, Complex structures that even partially extend into the ghosted region are duplicated in their entirety to the neighboring processor to ensure that each processor has access to all necessary information for handling bimolecular reactions. When assessing for steric overlap, only overlaps with complexes in the same processor and the ghosted complexes from the adjacent processor are considered. If a large complex overlaps with a non-ghosted complex from the neighboring processor, the neighboring processor will resolve this in the next timestep.

### 2.5. Synchronization of computation and communication

Adjacent processors cannot operate simultaneously to make decisions on the duplicated molecules in the ghosted regions, which interact with molecules on both processors. This interdependency requires synchronization of computation and communication to maintain accuracy (Fig 1D). We divide each processor’s region into left and right halves (Fig 1A). For the computation of 0^th^ and 1^st^ order reactions, which are not pairwise, the calculations proceed simultaneously and independently on each processor across the full region (Fig 1D step 1). Each processor then independently and simultaneously evaluates the binding probabilities of all pairwise reactions involving its own molecules and all ghosted molecules (step 2). Then, all processors update all molecules in their left half, including the left ghosted region, by deciding whether to perform reactions, and if so, performing those reactions. Otherwise, all molecules in the left half, *excluding* the ghosted region, perform diffusion events (step 3). Upon completion, each processor communicates information about all Molecules and Complexes in their left ghost region and their left edge region (which is the right ghost region for the neighbor), sending to their left processor (step 4). Processors update these molecules in their right half and right ghost region. They then proceed to decide all remaining pairwise reactions in their right half and perform selected reactions. Molecules in the right half can react with right edge molecules if they have not already reacted (step 5). Finally, the updated right edge and ghost regions are communicated back to the processor on the right (step 6). This staggered approach to computation and communication minimizes idle time, as computation proceeds simultaneously on all processors as much as possible. This strategy requires that the largest complex should not exceed half of the processor’s region, as larger complexes can straddle the border of the left and right halves simultaneously, which is not supported by the current protocol. We note here that the initial two steps could also be trivially accelerated using shared-memory threading via OpenMP, which is easy to implement for parallelizing loops over independent computations^51^. However, the subsequent steps require decision making on performing reaction or diffusion events that are not independent and thus not readily multi-threaded. This approach ultimately did not provide good scaling even for a few CPUs for NERDSS. We provide pseudo-code in the SI.

### 2.6. Communication load and structure

We implemented serialization and deserialization methods to transfer Molecule and Complex structures between processors. The serialization step packages the two data structures into a single MPI_CHAR array, enabling efficient transmission via non-blocking MPI functions MPI_Isend and MPI_Irecv^18^. In serial NERDSS, each molecule and complex object is assigned an index that corresponds to its position in the respective vector data structures. In parallel NERDSS, each molecule and complex object also needs a unique global identifier (ID) that remains unchanged throughout the object’s lifetime, even as the local index can vary. This global ID is needed to identify and track objects across different processors, and is used during the serialization and deserialization processes to correctly interpret and reconstruct the relationships between objects across processor boundaries. The deserialization process receives the MPI_CHAR using MPI_Irecv function, unpacks the two data structures from the MPI_CHAR variable, and reverses this mapping, converting the global IDs back to the appropriate local indices within the receiving processor’s local data structures. The MPI_Wait function ensures all necessary data has been received and processed before proceeding with the next computation step. After the deserialization process, the molecules and complexes that were sent but not received back are removed from the sending processor, as they have moved out of the edge or ghosted regions.

### 2.7. Analyzing scalability in parallel NERDSS

To identify the factors controlling scalability in parallel NERDSS, we differentiate between computation time (*T*_comp_) and communication time (*T*_comm_) (Fig 1D). *T*_comp_ represents the time spent performing the actual computations throughout the simulation iterations, such as executing reaction and diffusion events, and excludes the overhead of setting up the simulation, *T_setup_*. *T*_comm_ encompasses the time spent on serializing molecules and complexes, sending and receiving data using MPI functions, and deserializing the received data back into molecule and complex objects. As the number of processors increases, *T*_comp_ is expected to decrease linearly, assuming an even distribution of the computational workload among the processors, which is what we find for all systems. For our method, we expect *T*_comm_to decrease slightly with increasing processors, because with more processors, the domain size per processor shrinks, reducing the number of molecules that need to be looped through during serialization. We average *T*_comp_ and *T*_comm_ across all processors.

We quantify strong and weak scaling of the parallel code with an increasing number of processors, within and across nodes. For strong scaling, the problem size is fixed and thus the per-processor region shrinks with increasing processors. We compare the total simulation time on *p* processors, 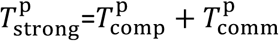 to the time on 1 processor. The speedup is defined as

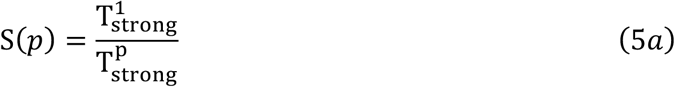

And the efficiency as:

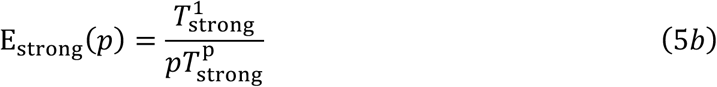

Linear scaling produces S(*p*) = *p*, and E_strong_(*p*) = 1. For the weak scaling, we keep the region size (and thus workload) per processor fixed and increase the total system size with each added processor. In this case, the efficiency is now measured as

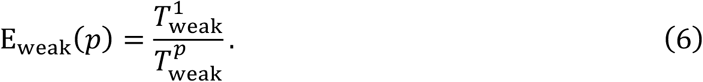

Deviations in both strong and weak scaling from ideal speedups and efficiency eventually accumulate due to increased communication overhead relative to computation time, data distribution inefficiencies, and/or memory bandwidth limitations.

### 2.8. Benchmarking systems for parallel NERDSS

#### Hardware and software used for benchmark

Benchmarking tests for evaluating the performance of parallel NERDSS were conducted on the Rockfish cluster using nodes equipped with the following hardware: (i) CPU: Intel Xeon Gold Cascade Lake 6248R, (ii) RAM: 192GB DDR4 2933MHz per node, (iii) Cores: 48 cores per node, (iv) The Rockfish cluster has Mellanox Infiniband High Data Rate 100 Gbps (HDR100) connectivity (1:1.5 topology). The following software versions were employed in the benchmarking process: (i) Open MPI: Version 4.1.1, (ii) C++ Compiler: GCC 9.3.0, (iii) GNU Scientific Library (GSL): Version 2.7.

#### Bimolecular reaction system

As a unit test, we simulate a reversible bimolecular reaction A + B ⇌ A.B where all species start off as monomers. We evaluate four different reaction environments (Table 1): (1) 3D; (2) 2D; (3) From 3D to 2D (explicit 2D molecule), where one reactant is in 3D and the other is constrained to 2D (4) From 3D to implicit 2D molecules represented by a density field. We run three independent trajectories per system and report the mean and standard error of the mean (SEM) of the species concentrations in time.

**Table 1.** System and reaction parameters for the bimolecular, trimer, and hexamer systems. All bimolecular systems use box dimensions of [20000, 2000, 2000] nm.

| Reaction | System details | Initial # | Diffusion<br>Constant ($D_x$,<br>$D_y$, $D_z$), $\mu\text{m}^2/\text{s}$ | Forward<br>Rate ($\mu\text{M}^{-1}$<br>$\text{s}^{-1}$) | Backward<br>Rate ($\text{s}^{-1}$) |
| --- | --- | --- | --- | --- | --- |
| A + B ⇌ A.B | A (3D) | 10000 | (10, 10, 10) | 100 | 10 |
|  | B (3D) | 10000 | (10, 10, 10) |  |  |
| C + D ⇌ C.D | C (2D) | 10000 | (2, 2, 0) | 0.53nm<sup>2</sup><br>/μs | 15.9 |
|  | D (2D) | 10000 | (2, 2, 0) |  |  |
| E + F ⇌ E.F | E (3D) | 10000 | (10, 10, 10) | 55 | 5.5 |
|  | F (2D) | 10000 | (2, 2, 0) |  |  |
| G + H ⇌ G.H | G (3D) | 10000 | (10, 10, 10) | 55 | 5.5 |
|  | H (implicit 2D) | 10000 | (2, 2, 0) |  |  |
| trimer: V=<br>[20000,2000,2000] | A (3D) | 20000 | (10,10,10) | 38.96 | 6.47 |
| trimer: V=<br>[5000, 100, 100] | A (3D) | 6022 | (10,10,10) | 10.86 | 901.6 |
| hexamer: V=<br>[20000,2000,2000] | A (3D) | 20000 | (10,10,10) | 43.08 | 7.15 |
| hexamer: V=<br>[5000,100,100] | A (3D) | 6022 | (10,10,10) | 11.16 | 926.3 |

For the strong scaling tests, the system’s geometry consists of a box with dimensions x, y, z = [20000, 2000, 2000] nm and 10000 initial molecules for each molecular type. In the weak scaling tests, the base system for the single-processor simulation is [2000, 2000, 2000] nm with 1000 initial molecules for each type. With increasing processors *p*, the box dimensions become [2000*p*, 2000, 2000] nm, and the initial number of molecules for each monomer type is set to 1000*p*. A membrane surface is located at the bottom of the z-axis. Molecules on the 2D surface are constrained to the membrane and can be either explicit diffusing particles or implicit molecules represented by a density field. The initial copy number and diffusion constant for each molecule in the general benchmarking system are listed in Table 1. Δ*t*=0.1 μs. Note the 2D system in Table 1 lists the 2D intrinsic rate, 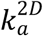. The dissociation also lists the intrinsic unbinding rate 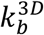. To determine the macroscopic rates used in the well-mixed rate equations, we calculate it using^21^ 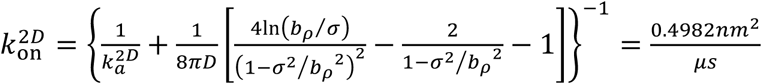, where 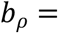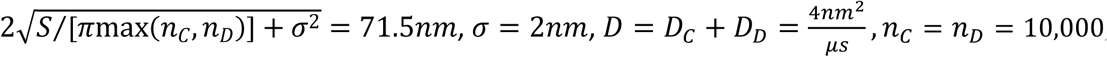, S = 40*μ*m^2^ and The macroscopic off-rate is: 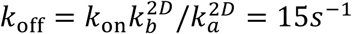.

#### Oligomer macromolecule assembly system

We benchmarked the higher-order assembly of a homo-trimer and a homo-hexamer, which form loops of 3 or 6 monomer subunits, respectively, along with all intermediates. Growth is not restricted to monomer addition, meaning for the hexamer, two trimers can combine into the hexameric loop. In both cases, each monomer subunit has two interfaces, labeled c and q, and they bind to each other only (e.g., c does not bind to c). This is encoded in the same reaction network for both systems: A(c) + A(q) <-> A(c!1).A(q!1). Each binding reaction only forms one bond. When the final subunit is added to the structure, it thus contains either 3 or 6 subunits, but with only 2 or 5 bonds. The formation of the final bond is no longer a bimolecular reaction, as the subunits do not diffuse relative to one another. The binding is evaluated as a 1^st^ order reaction or loop-closure reaction, and the rate is defined by the relation *k*_close_ = *k*_f_c_0_exp(−Δ*G*_coop_/*k*_B_*T*), where *c*_0_ = 1*M* and here we have no cooperativity, Δ*G*_coop_ = 0, meaning the free energy of the final bond is the same as all other bonds. Δ*t*=0.1 μs. For monomers, *D*_=_=10 μm^2^s^−1^ and *D*_rot’_= 0.1 rad^2^μs^−1^. The key difference between the trimer and hexamer is the orientation that the monomers adopt upon binding: the trimer forms 60 degree angles between a monomer center and its two binding partner centers, and the hexamer forms 120 degree angles between the same. The binding radii are slightly different, *σ* = 0.73nm and 1nm for the trimer and hexamer, respectively. Reaction parameters are in Table 1.

#### Protein lattice formation in 3D or 2D

The formation of a flat hexameric lattice sheet evolves from the self-assembly of triskelia-shaped monomers inspired by the clathrin triskelia^32^. Each triskelion has three interfaces to bind with other triskelia and three additional interfaces that bind to another protein component, AP. Each AP monomer has two interfaces, one to bind the triskelia and one to bind the membrane lipids. We initiated the system with 10,000 triskelia, 10,000 AP, and 204,082 implicit lipids. The volume is [20000, 2000, 2000] nm and Δ*t*=0.1 μs. For transport coefficients, *D*_t,trisk_=13 μm^2^s^−1^ and *D*_rot, trisk_= 0.03 rad^2^μs^−1^, *D*_t*, AP*_ =25 μm^2^s^−1^ and *D*_rot,AP_= 0.5 rad^2^μs^−1^, *D_t,lipid_*=0.5 μm^2^s^−1^ and *D*_rot,lipid_= 0.01 rad^2^μs^−1^. All reactions can occur in both 3D and 2D. The 2D rates are distinct, and we define them relative to the 3D rates by 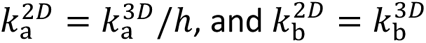. These are the intrinsic rates used for the reaction probabilities (Eq. 1). Intrinsic rates in 3D convert between macroscopic 3D rates using 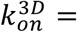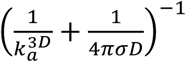. In 2D, there is no single macroscopic rate, so we only define 2D intrinsic rates relative to 3D intrinsic rates^21^. The reaction network is 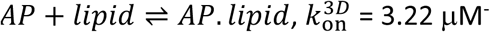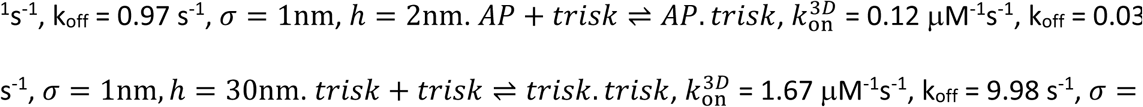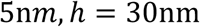. When the triskelia binds to AP, its association rate to itself increase 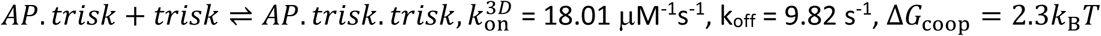. This model and parameters were optimized in previous work^52^, but we accelerate association rates here to nucleate lattices at the distinct concentrations used here. We ran 8 independent trajectories per system.

#### Comparison with theory and continuum results

The ordinary differential equations (ODE) for the trimer assembly model non-spatial kinetics were solved using Python’s scipy.integrate.solve_ivp() for numerical integration (see SI for model definition). The hexamer assembly thermodynamic equilibrium calculation was converged iteratively using MATLAB code. The partial differential equation (PDE) solution for the 3D to 2D binding were solved using Virtual Cell (version 7)^5^, employing a fully implicit finite-volume method on a regular grid (variable timestep, max timestep of 0.1s), with absolute error: 10^−9^, and relative error: 10^−7^. The mesh side length is ∼0.022μm. The bimolecular association kinetics for well-mixed reactants in 3D and 2D uses the analytical solution.

## 3. RESULTS

### 3.1. Accuracy and scaling of 3D bimolecular reaction benchmark

Reversible bimolecular reactions (A + B ⇌A.B) represent a foundational unit test of the reaction-diffusion software, dependent on diffusion of particles to collision and evaluation of binding/unbinding between all A, B reaction pairs. For both the strong scaling and weak scaling systems illustrated in Fig 2, our parallel code across *p*=1 to 96 processors produced excellent agreement with the theoretical kinetics and equilibrium expected from well-mixed reactants, as expected given our well-mixed system initialization (Fig 2C and 2E). Additional validation at varying binding constants (Fig S2) also exhibits excellent agreement with kinetics derived from solving the mass-action rate equations. From these simulations we also verified the mean-squared displacement (MSD) of the particles correctly obeys the Einstein relation. We evaluated displacement in the x-dimension, to minimize the effects of the reflecting boundaries, and compare with 〈(*x*(*t* + Δ*t*) − *x*(*t*))^2^〉 = 2*D*Δ*t* in Fig 2D, showing excellent agreement until the longest Δ*t*, where the simulated MSD shows evidence of boundary effects. Because the diffusion constants of our dimers is *D_dimer_* = *D*_c_/2, the diffusivity of the population slows until equilibrium is reached. We plot the resulting MSD of A molecules at equilibrium where 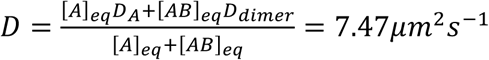

**Fig 2.**
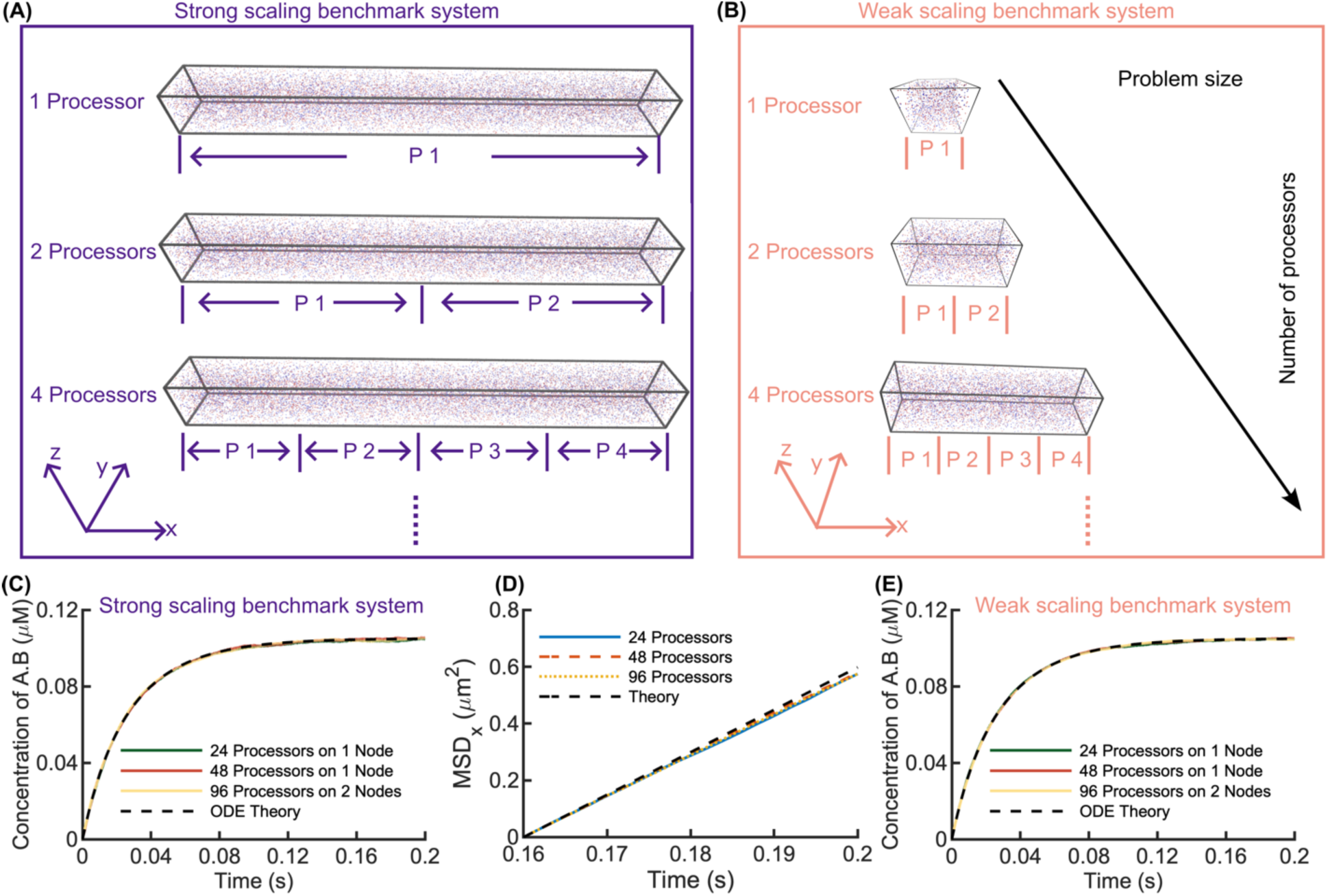
Validation of 3D bimolecular reaction system (A+B⇌A.B) accuracy using parallel NERDSS. (A) Schematic of the strong scaling benchmark system, where the simulation system size remains constant across varying the number of processors . (B) Schematic of the weak scaling benchmark system, where the simulation box size and molecules copy numbers increase linearly with the number of processors. (C) Concentration of the product A.B over time for the strong scaling system shows excellent agreement with the well-mixed solution to mass-action kinetics (black dashed), which matches our system conditions. Simulated with 24, 48, and 96 processors. (D) Mean square displacement (MSD) vs. time for different processors counts in the strong scaling system, validating diffusion accuracy in parallel NERDSS. The black dashed line represents the theoretical MSD in the x dimension. (E) A.B concentration over time for the weak scaling system is equally accurate. All simulation results are presented as mean ± standard error of the mean (SEM), which here is smaller than the linewidth. See Table 1 for reaction parameters. The R_cut_ is 12.4 nm. The total system is divided into [1613, 4, 4] sub-volumes with each sub-volume of size [12.4, 500, 500] nm.

We assess the scalability of our parallel code from *p=*1 to 96 processors, comparing the speed also across *n*=1-8 nodes (Fig 3). The computation time for the strong scaling system decreases linearly as expected, showing ideal speed ups for computation (Fig 3A). We do not see any speed loss by distributing the computation across multiple nodes, in fact both the computation and communication times are slightly reduced by using more nodes with fewer processors per node, likely due to increased RAM (Fig 3B). Efficiency remains close to 100% up to *p*=8 and ∼90% at *p=*24 but drops to 63% at 96 processors as computation (501 seconds / 2 × 10^6^ iterations) and communication (428 seconds / 2 × 10^6^ iterations) time become comparable (Fig 3A, C). Our best scaling results (>80x increase with 96 processors) are shown for systems in Fig 4. The communication time for all systems is strongly influenced by the size of the ghosted region (Fig 1C), which we set to be its minimum size of *R*_cut_(Eq. 3), dependent on Δ*t* and diffusion constants. The communication load remains the same as more processors are added, given the constant size of *R*_cut_. However, the communication time decreases slightly because we loop over all molecules per region prior to communication to identify molecule IDs for passing between processors, and the region size shrinks with *p*.

**Fig 3.**
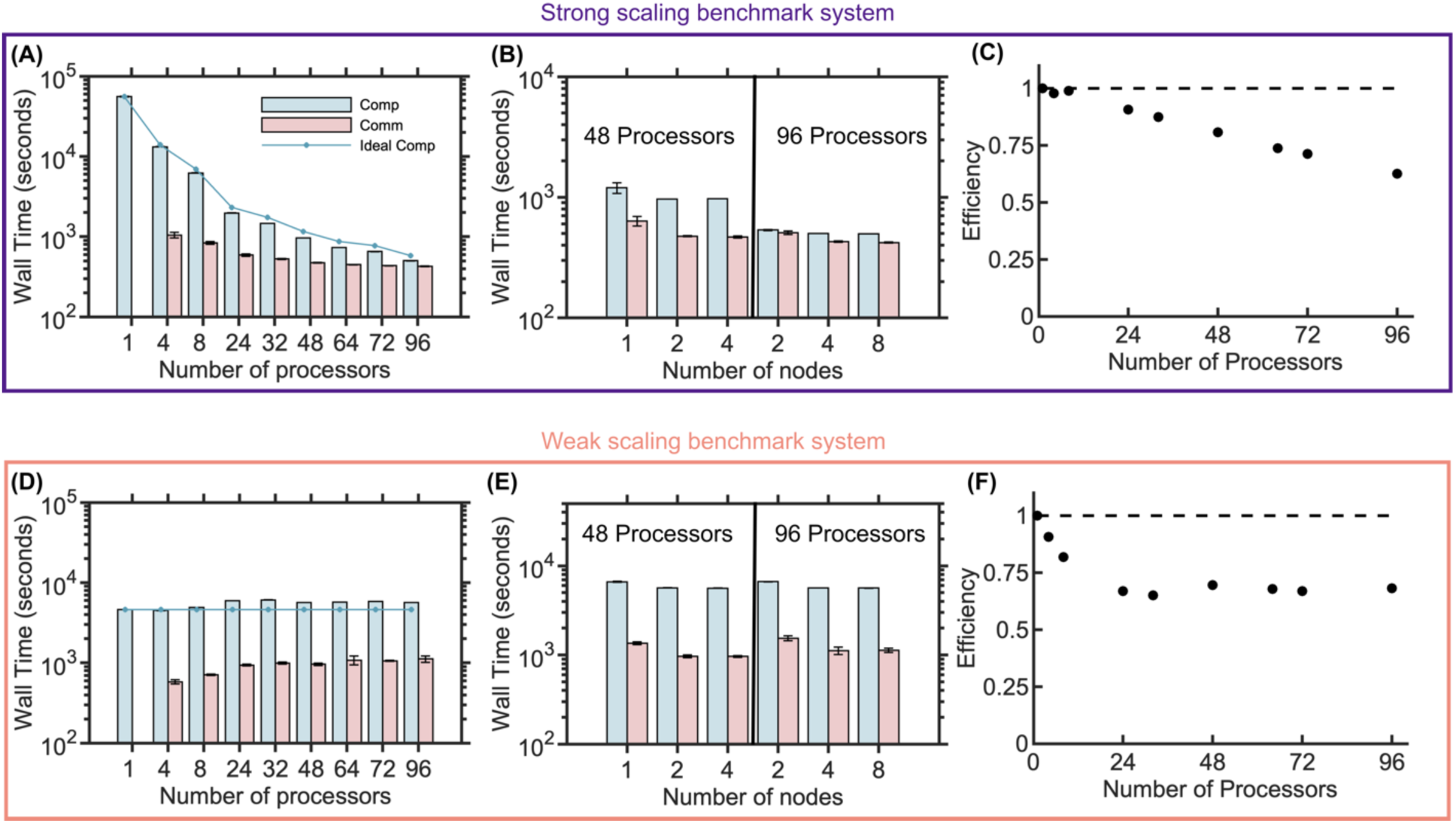
Strong and weak scaling benchmarks of parallel NERDSS for 3D bimolecular reactions show accelerated simulations up to 96 processors. (A) Computation and communication times for strong scaling across various processor counts. Bars indicate times measured from the simulations along with standard deviation across 3 trajectories. The connected points track an ideal linear speed-up in computation time only, showing that our computation does scale linearly across processors. The wall time is measured over 2 × 10^6^ iterations (0.2s) for all simulations. (B) Computation and communication times for strong scaling using 48 processors or 96 processors distributed across multiple nodes shows more nodes can improve speed for the same number of processors. (C) Strong scaling efficiency as a function of the number of processors. (D) Computation and communication times for weak scaling with increasing numbers of processors, or with (E) either 48 processors or 96 processors but distributed across multiple nodes. (F) Weak scaling efficiency as a function of the number of processors. For (A) and (D) 1 node was used for 4 to 32 processors, 2 nodes for 48 to 72 processors, and 4 nodes for 96 processors.

**Fig 4.**
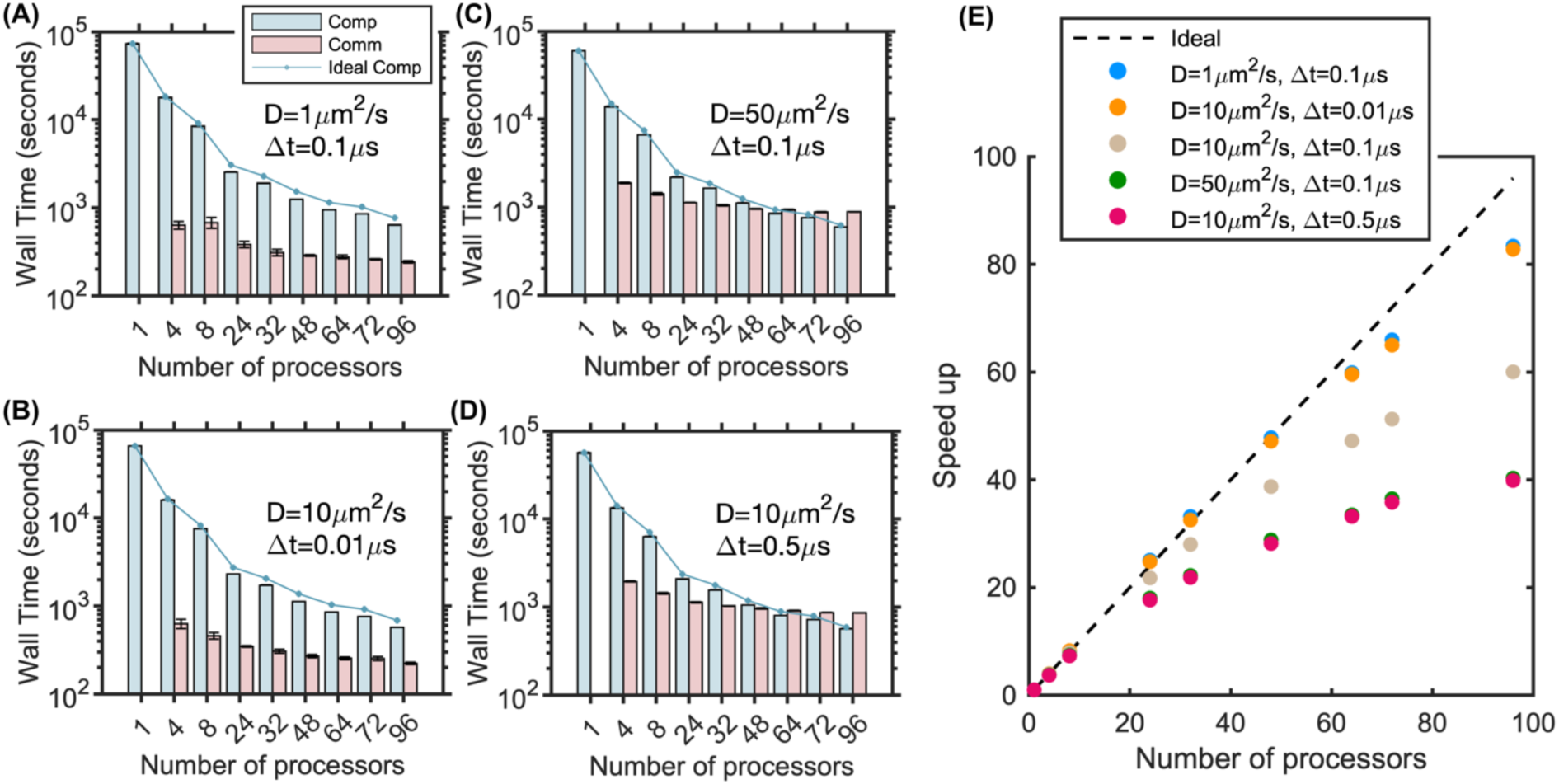
Strong scaling benchmark of parallel NERDSS for 3D bimolecular reactions shows optimal scaling for slow diffusion and/or short timesteps. Computation and communication times for (A) D=1μm2/s, ∆t=0.1μs (blue dots). (B) D=10μm2/s, ∆t=0.01μs (orange dots). (C) D=50μm2/s, ∆t=0.1μs (green dots). (D) D=10μm2/s, ∆t=0.5μs (red dots). Points indicate the ideal computation time for linear scaling (Eq. 2). Wall time is reported over 2 × 10^6^ iterations for all simulations. (E) Speedup vs. processor count for all cases. The dashed line represents ideal speedup (Speedup = n), illustrating the best performance with slower diffusion or shorter timesteps (blue and orange dots). 1 node was used for 4 to 32 processors, 2 nodes for 48 to 72 processors, and 4 nodes for 96 processors. In the simulations for C and D, 1 node was used for 4 to 8 processors, 2 nodes for 24 processors, 3 nodes for 32 processors, 4 nodes for 48 processors, 6 nodes for 64 to 72 processors, and 8 nodes for 96 processors.

For the weak scaling benchmark, the computation and communication times both increase slightly up to *p*=24, and after that remain steady, indicating good performance for large systems (Fig 3D). Efficiency drops to 67% for 24 processors and then remains around 68% as the processor count increases further to 96. Again, we see that splitting processors across nodes provides a small speedup, indicating communication across nodes is not rate limiting (Fig 3E), and we find the scalability improves for a benchmark with half the size and copy numbers (Fig S4). Finally, we note the overhead set-up time for the simulations does increase slightly for the strong scaling system as *p* increases but remains under 200s (Fig S3). For the weak scaling system, for the largest systems the set-up time increases significantly (Fig S3), indicating that additional optimization in the set-up could be improved for the largest problems.

We systematically tested the role of the length scale *R*_cut_ (see Eqs 2, 3) on the strong scaling of the bimolecular benchmark by keeping the copy numbers and system size the same, while varying diffusion constants *D* and timesteps ∆t (Fig 4A-D). As expected, decreasing either ∆t or *D* decreased the size of *R*_cut_ and thus caused an improvement in scaling and parallel efficiency, allowing us to achieve our best scaling results of >80 fold speed-ups on 96 processors (Fig 4A, 4B, 4E). These 10-fold changes reduced *R*_cut_ from 12nm to 5.3nm, reducing the communication load. Conversely, increasing the diffusion constant by a factor of 5 (Fig 4C) or increasing the timestep by a factor of 5 (Fig 4D) raises R_cut_ to 25.2nm, leading to a reduction in scaling and parallel efficiency (Fig 4E). Here again we took advantage of slightly better performance for systems with larger communication loads by using more nodes and thus fewer processors per node.

Particle density only impacts the scaling slightly, with higher particle density (System 5) resulting in slightly worse scaling as we keep the volume and geometry fixed (Fig 5). The communication cost increases nonlinearly with the number of particles, in part due to the serialization steps that require all particles per processor are looped over, a step that in future work could be optimized (Fig 5C). Lastly, because our parallel NERDSS is partitioned across processors only in the x-dimension, the scalability is sensitive to the geometry of the volume. In Fig 5, all systems have the same volume, but 3 systems vary in the length of x relative to y and z. Not surprisingly, the systems with the largest asymmetry in the x-dimension showed the best scaling, as this results in a reduced y-z interface and therefore fewer total particles at the processor boundaries, significantly reducing communication load between processors. A cubic volume produces the worst scaling (System 1-beige dots), whereas the system with the largest asymmetry ([40000,1414,1414] nm) produces nearly perfect scaling up to 32 processors (System 3-green dots).

**Fig 5.**
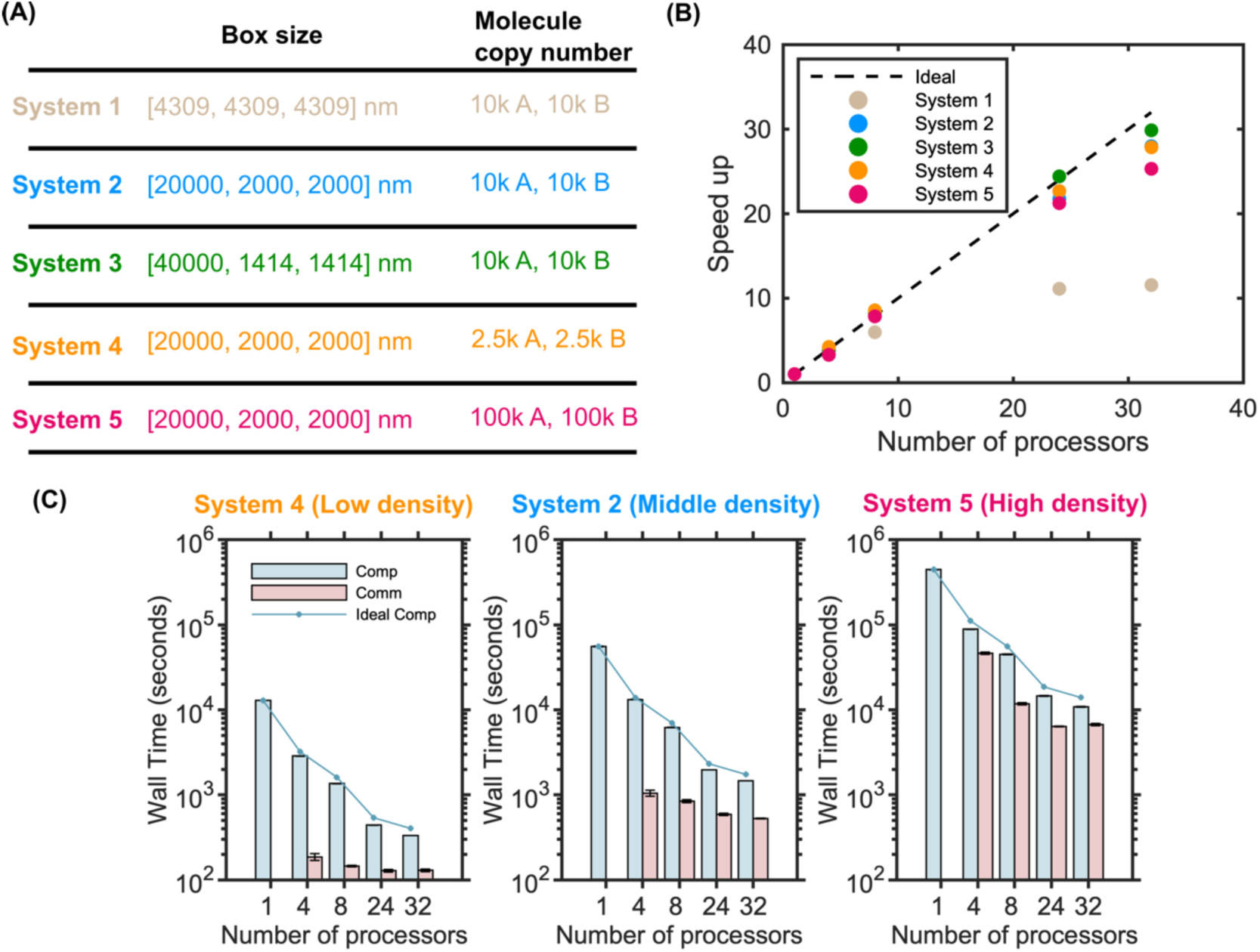
Increased molecular density and asymmetry in the x-dimension improves strong scaling performance for 3D bimolecular reactions. (A) Box size and molecular copy numbers of each system configuration. All systems have the same total volume. (B) Speedup vs. processor count for each system. Increasing the asymmetry along the x-axis results in improved scaling with the same volume and particle numbers, as seen by comparing System 1 (beige) to System 2 (blue) to System 3 (green). Higher particle density results in a slight performance drop in the scaling, as seen by comparing System 4 (orange) to 2 (blue) to 5 (red). (C) Computation and communication times for increasing particle densities from System 2, 4, and 5, showing a more dramatic increase in communication time relative to computation time for the highest densities. Wall time is reported over 2 × 10^6^ iterations for all simulations. All results collected within a single node.

### 3.2. Accuracy and scaling of bimolecular benchmarks in 2D and 3D to 2D

We validated the accuracy of parallel NERDSS for bimolecular reactions purely in 2D and for reactions where one partner is in 3D and the other is restricted to the 2D surface, showing excellent agreement with expected kinetics from continuum results (Fig 6). The kinetics of the 3D-to-2D systems are not well described by a well-mixed ODE model because diffusion to the surface slows the overall kinetics, thus requiring comparison to a PDE model. The system volume is the same as for the initial 3D benchmarks (Fig 2), but now the membrane/2D surface is defined as the x-y plane at z=0. The 2D molecules are treated as explicit particles (Fig 6A2) and as an implicit density field (Fig 6A3), which is a faster calculation as there is no propagation of 2D particles and only one pairwise evaluation per 3D molecule with the surface. For the 2D system, the parallel efficiency for both strong scaling and weak scaling remains 100% up to 24 processors, demonstrating the excellent scalability of parallel NERDSS for 2D bimolecular reactions (Fig 6B1,C1, D1). For the explicit particles 3D to 2D simulations, the parallel efficiency for strong scaling remains above 96% up to 32 processors Fig 6(B2, C2, D2) and for weak scaling remains above 68% up to 96 processors. Finally, for the bimolecular reaction from 3D to implicit 2D, the system has significantly better weak scaling performance compared to the 3D to explicit 2D system, driven by the decreased communication load resulting from the implicit representation of the 2D particle density.

**Fig 6.**
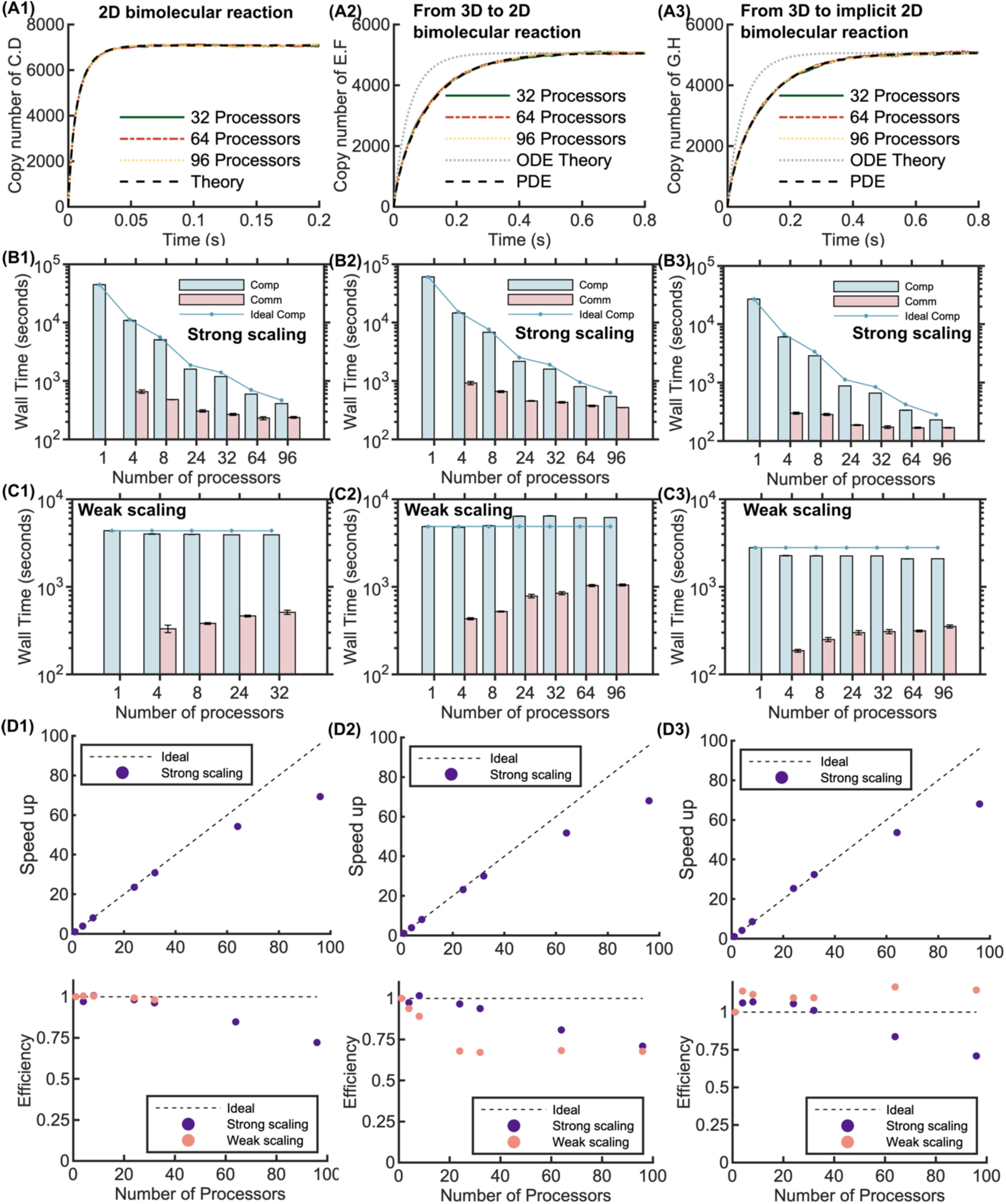
Accuracy and scaling for bimolecular reactions on 2D, from 3D to 2D, and from 3D to implicit 2D using parallel NERDSS. (A1-A3) The three different bimolecular reaction environments produce excellent agreement with solutions to continuum differential equations (black dashed). The left column is 2D results, the middle column is 3D to explicit 2D particles, and the right column is 3D to implicit 2D particles. For the 3D to 2D systems, we compare the NERDSS results to the time-dependent solution using continuum PDEs (black dashed), as diffusion to the surface produces a delay in the association kinetics compared to a well-mixed system (gray dashed). For the 2D reaction system, *R*_cut_ = 5.8 nm, and the strong scaling system is divided into [3451, 2, 2] sub-volumes. 1 node was used for 4 to 32 processors, 6 nodes for 64 processors, and 8 nodes for 96 processors. For both bimolecular reactions from 3D to 2D, *R*_cut_= 9.8 nm, and the strong scaling system is divided into [2036, 3, 3] sub-volumes. 1 node was used for 4 to 8 processors, 2 nodes for 24 processors, 3 nodes for 32 processors, 6 nodes for 64 processors, and 8 nodes for 96 processors. (B1-B3) Computation and communication times with different numbers of processors for each environment on the strong scaling system, with linear scaling of computation time shown in connected dots. (C1-C3) Computation and communication times for the weak scaling benchmark for a 0.2-second simulation (2 × 10^6^ iterations). (D1-D3) Total speedup of the strong scaling (upper panel) and the efficiency of strong and weak scaling (lower panel) for each environment.

### 3.3. Accuracy and scaling of oligomer assembly systems

In Fig 7, we demonstrate the accuracy and scaling of higher-order assembly in parallel NERDSS via the formation of trimers and hexamers, each assembled from one type of monomer subunit. The kinetics of monomer assembly into dimers and trimers from parallel NERDSS shows excellent agreement to the well-mixed results from solving the corresponding system of ODEs (Fig 7A1). We expect good agreement between the kinetics of our spatial stochastic simulations and the ODEs because our monomers are well-mixed, and the reaction dynamics are not diffusion-limited. The accuracy of the hexamer assembly is validated against the equilibrium steady-state (Fig 7A2). Both systems show good scaling up to 36 processors (Fig 7B-C), with a loss of efficiency in speed-up due to the increased communication costs, which are also higher for the hexamer. The hexamer system has a higher *R*_cut_(24.57nm) than the trimer system (*R*_cut_ = 14.51*nm*) due to the different *σ*, which significantly increases the communication load. Further, because any molecule in the ghosted region must be communicated to the neighboring processor along with the complex it belongs to, larger complexes like the hexamer increase the size of the communicated data structures.

**Fig 7.**
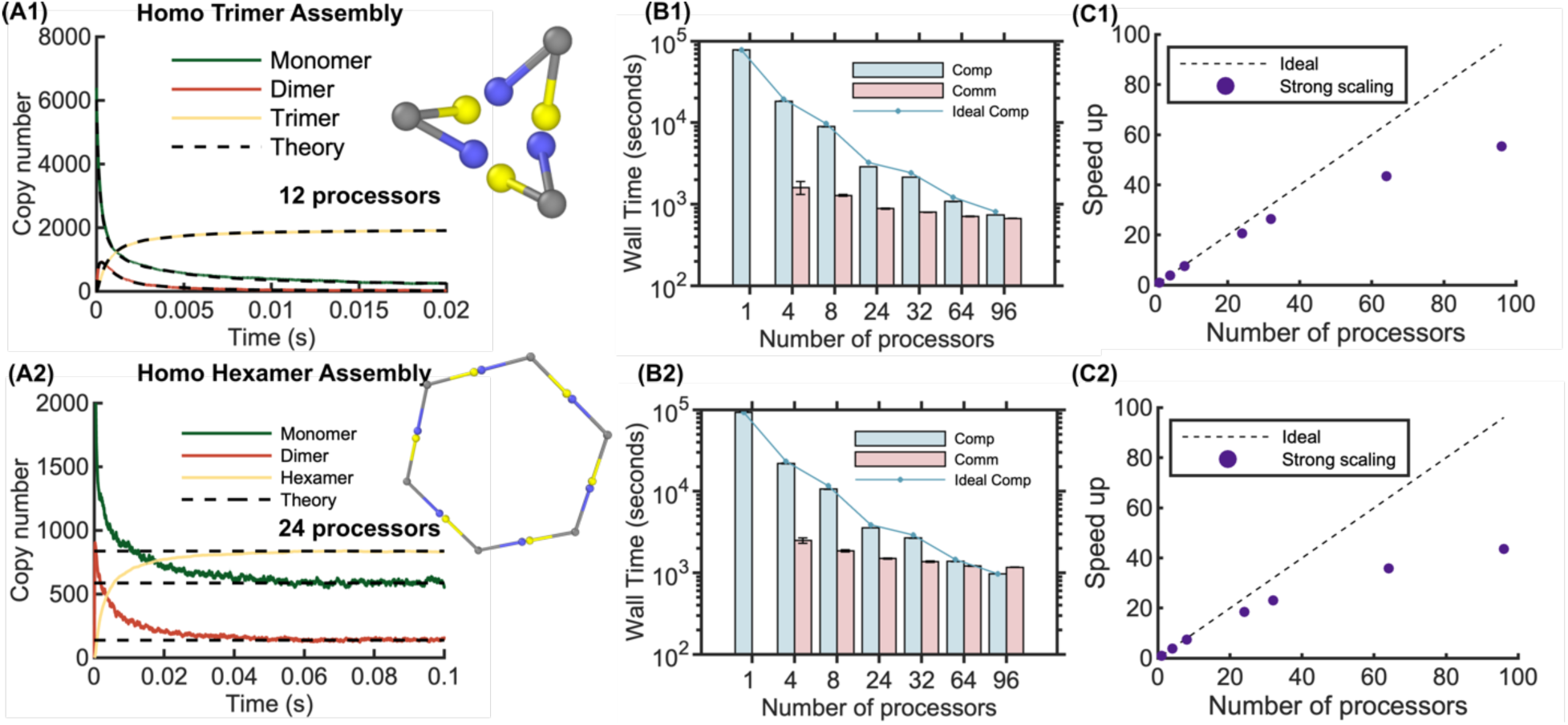
Benchmarking results for macromolecule assembly of trimers and hexamers using parallel NERDSS. Top row shows the accuracy and scaling results for monomers assembling into a trimer. All monomers are the same and have two unique sites, c (yellow) and q (blue) that only bind each other. (A1) The assembly of monomers (A, green) into dimers (A_2_, red) and then trimers (A_3_, yellow) from NERDSS parallel on *p*=12 shows excellent agreement to the well-mixed kinetics computed from the corresponding system of ODEs (black dashed). Reaction parameters in Table 1 for trimer in box dimensions [5000, 100, 100] nm. (B1) Compute and communication times for the strong scaling on the trimer in box dimensions [20000, 2000, 2000] nm, reaction parameters in Table 1. (C1) Speedups for the strong scaling. 1 node was used for 4 to 8 processors, 2 nodes for 24 processors, 3 nodes for 32 processors, 6 nodes for 64 processors, and 8 nodes for 96 processors. Bottom row shows accuracy and scaling results for assembly of hexamers. (A2) The equilibrium population of hexamers, monomers, and intermediates reached using parallel NERDSS on 24 processors shows excellent agreement with the calculation from equilibrium thermodynamics. Geometry shown in inset. Reaction parameters in Table 1 for trimer in box dimensions [5000, 100, 100] nm. (B2) Timings for the strong scaling benchmark on the hexamer in box dimensions [20000, 2000, 2000] nm, reaction parameters in Table 1. (C2) Speed ups for the strong scaling benchmark. All times reported for 2 × 10^6^ iterations (0.2 s).

### 3.4. Benchmarking lattice sheet formation from 3D to 2D

Our final benchmark illustrates the accuracy and scaling of parallel NERDSS for a more complex system with multiple distinct components driving nucleation of extended hexagonal lattices on the membrane surface. This type of model dynamics cannot be captured with continuum methods or predicted from analytical theory, and thus we validate the kinetics of component binding relative to the serial version of NERDSS (Fig 8A). The triskelia-shaped monomers were designed to study the mechanisms of clathrin lattice nucleation and growth in cell biology^32^. The triskelia bind each other and another protein that helps facilitate their assembly and more importantly, localizes them to the 2D surface where they exploit dimensional reduction, or the reduction in search space for diffusion-driven binding^53^. Hence most of the assembly occurs on the 2D surface^54,55^. This system has good scaling up to 20 processors, but suffers at higher processor numbers due to the higher communication costs. Here again, the formation of the larger assemblies is driving up communication costs. Below we discuss strategies for future improvement of parallel NERDSS in more efficiently dealing with the largest assemblies.

**Fig 8.**
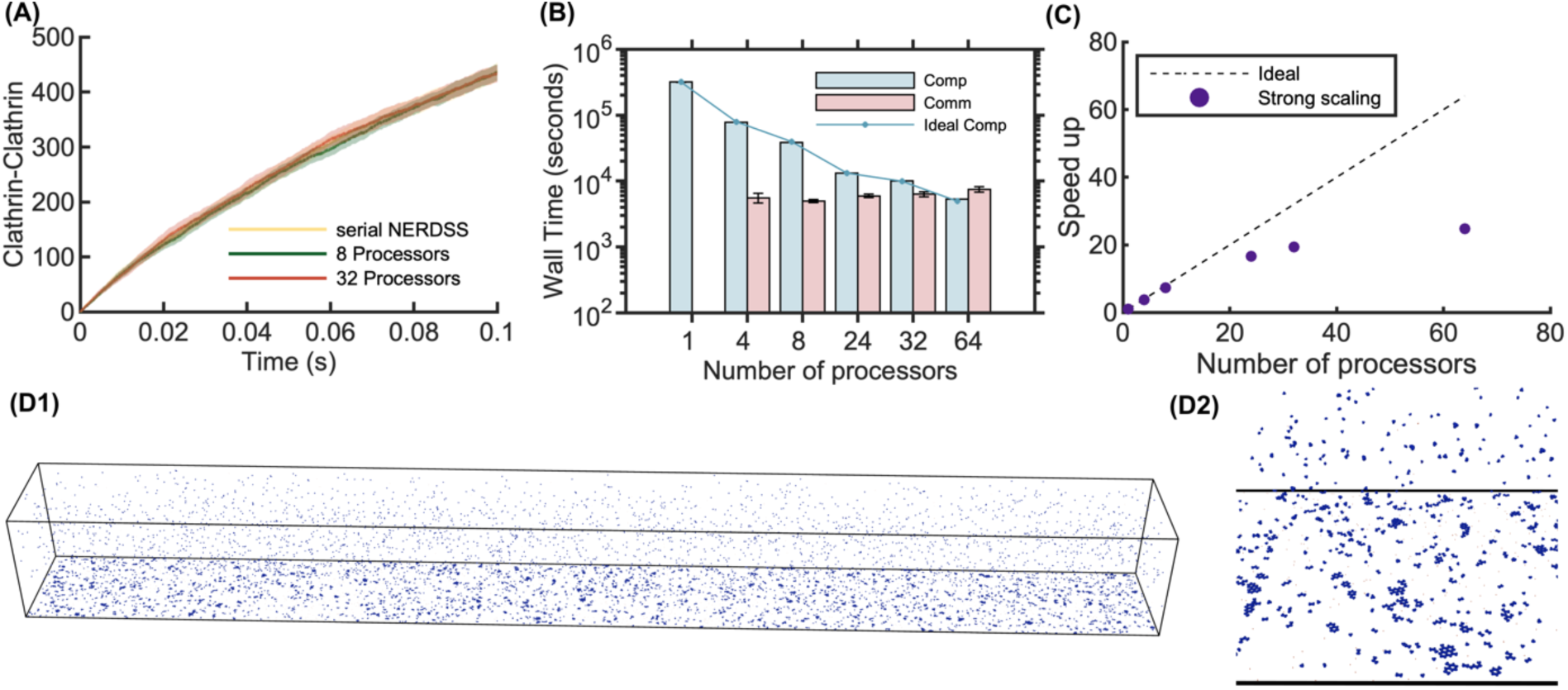
Benchmarking results for lattice self-assembly using parallel NERDSS. (A) Time dependence of the number of triskelia-triskelia bonds formed. (B) Computation and communication times vs processor numbers for strong scaling, with the dotted line indicating linear scaling for compute time. (C) The corresponding strong scaling speedup. Times reported for 2 × 10^6^ iterations (0.2s). (D1) A simulation snapshot running across 32 processors at 2s. (D2) A magnified view of a selected region from the snapshot showing the formation of hexagonal lattices on the 2D surface only.

## 4. DISCUSSION and CONCLUSIONS

Particle-based reaction-diffusion software are unique amongst reaction-diffusion solvers for their ability to capture the volume-excluding, discrete size, and higher-order structural assembly of component species that fundamentally control dynamics in many chemical, biological, and engineered systems. Our MPI parallelization of the particle and rigid-body reaction-diffusion software, NERDSS, demonstrates that we can achieve up to nearly 90-fold speedups on 96 CPUs for reversible bimolecular association in various spatial environments (3D, 2D, transitioning between), enabling the simulation of large systems over tens of microns while maintaining the nanometer-scale resolution of molecular components. The parallelization using MPI enables distributed computing across multiple nodes as well as within single nodes; the scaling even improves slightly when more nodes with fewer processors per node are used. The excellent scaling for smaller numbers of processors (up to 20) across all our benchmark systems will also facilitate improved parameter estimation techniques, where fast simulations are critical for effective ‘learning’ using a variety of optimization techniques^56,57^. Our benchmarking from simple bimolecular reactions to complex macromolecular assemblies and large-scale lattice formation that can only be simulated with particle-based methods like NERDSS, demonstrates that the accuracy of the governing reaction dynamics is retained with the enhanced performance of parallel NERDSS.

Our analysis on the parallel scaling and efficiency as it depends on the particle density, system geometry, and timestep parameters illustrates which systems have the best performance while also highlighting areas for improvement over our current implementation. In general, the scaling across >20 processors suffers with any change to system parameters that increases the communication load by increasing the ghosted domain length *R*_cut_in the *x* dimension or its interface across nodes (*boxl*_z_ · *boxl*_y_). To better deal with any system geometry, the parallelization can be extended to work across the *z* and *y* dimensions in the same way as across the *x* dimension, as successfully implemented by MD software^38,39,44^.

However, a more important step would be to improve handling of the higher-order assembly systems, as we found these systems did not scale as well as the bimolecular systems. Like the bimolecular association, these systems did show linear scaling in the computation time, so increased communication is responsible for the observed decrease in parallel efficiency. This stems from the rigid-body dynamics of the assemblies. Although each molecule makes its own decisions about reactions to perform based on nearby partners, the position updates are coupled for all molecules within a rigid-body complex. While MD also deals with long-range coupling, it performs a single position update for each particle on its home processor. In NERDSS, because reactive partners must avoid overlap, position updates at times must be iterated on for a single step, which currently requires a larger set of molecules beyond the ghosted domain are shared across processors. We could employ more sophisticated strategies for hierarchical updates of molecules, prioritizing displacement of larger, slow-moving assemblies first to reduce required data sharing while still ensuring smaller partners exclude volume. Our data structure management for the ghosted regions could benefit from further optimization, as these molecules are embedded in the full data structure for the processor, increasing the time searching within memory for the relevant data. Neighbor lists could also be more naturally incorporated into the identification of ghosted domain particles, as that step currently considers all molecules within the processor. These same strategies will be important for addressing a current limitation of parallel NERDSS: it allows formation of complexes up to half the processor domain size. While this is sufficient for many macromolecular complexes or lattice assemblies on the scale of <100nm, as shown above, both filaments and 2D lattice structures in certain parameter regimes can assemble to span longer distances. In addition to the steps indicated above, multi-level decomposition schemes used in MD simulations handle short-range Van der Waals forces and long-range electrostatic interactions efficiently^26^, and a similar strategy could be used in parallel NERDSS for handling extended assemblies.

The MPI parallelization strategy we adopted here should transfer readily to other particle-based reaction-diffusion software that are either implemented in serial or using GPUs, which do not typically scale across more than a single GPU^13,14^. To this end, we provide additional documentation in the SI that extends from the Methods section above to detail the necessary modifications to serial code for adding communication and handling ghosted and edge domains, while otherwise minimizing modifications to the majority of the serial C++ subroutines (SI). The code is available open-source under a GNU license at github.com/mjohn218/NERDSS/tree/mpi. Particle-based RD tools are an important complement to continuum RD approaches, having been applied to uniquely quantify mechanisms of self-assembly^16,30,31^, condensate formation^29^, localized signal transduction^23,58,59^, pattern formation^4,60^, crowding^61^, and membrane-mediated clustering^32,33,62^ with direct comparison to experimental data. Parallelization strategies such as ours are important steps to enhance the speed and thus tractability of these simulation tools.

## ACKNOWLEDGEMENTS

M.E.J. gratefully acknowledges funding from a National Institutes of Health MIRA Award R35GM133644, and XSEDE extended support via research allocation MCB150059. We acknowledge use of the ARCH supercomputer rockfish at Johns Hopkins.

