## Supplemental Information for "Parallelization of particle-based reaction-diffusion simulations using MPI"

For

### SUPPLEMENTAL METHODS

#### S1. Molecule and Complex data structures

The molecule serves as the fundamental simulation object within the system. Each molecule is characterized by a center-of-mass and one or more interfaces capable of binding to a single interface from another molecule. Multiple molecules can form a Complex via interface binding. A single molecule is also assigned to a Complex of size 1. Both intramolecular and intracomplex flexibility are not allowed; all species are rigid bodies.

The simulation environment is structured as follows:

1) A vector of molecules stores all molecules present in the system.
2) Each molecule is assigned a unique *index* property, corresponding to its position within the vector.
3) Each molecule contains a vector of interfaces.
4) Each interface is assigned an *index* property, representing its position within the molecule's interfaces vector.
5) Each interface can have more than one state. And the interface can change its state during the simulation.
6) A vector of complexes stores all complexes formed in the system.

7) Each complex is assigned a unique *index* property, corresponding to its position within the vector.
8) Each complex has a vector of integers named *memberList*, storing all indexes of molecules forming this complex.
9) Each molecule has a *myComIndex* property storing the index of complex to which it belongs.

This organizational structure allows for efficient tracking of molecular interactions and complex formations. An interface can identify its binding partner (both the molecule and the specific interface) by storing the corresponding indexes.

### S2. Sub-volume Optimization

To enhance computational efficiency, we previously implemented a sub-volume division strategy in the serial code. Each sub-volume only evaluates interactions within itself and to adjacent sub-volumes. In 3D, each sub-volume has 26 neighboring sub-volumes (ignoring boundaries).

1) *Cutoff Distance:* We calculate a cutoff distance based on the largest separation between species that can still result in a bimolecular reaction. Beyond this distance, molecules are too far apart to collide and interact with one another.
2) *Spatial Division:* The simulation volume is partitioned into sub-volumes, each with dimensions greater than or equal to the cutoff distance. This division guarantees that binding interactions between interfaces can only occur within the same sub-volume or between adjacent sub-volumes. This significantly reduces the number of interface pairs that need to be checked for potential reactions.
3) *Balance Consideration:* While increasing the number of sub-volumes improves efficiency, an excessive number will slow down the simulation due to the computational overhead of iterating through all sub-volumes and their adjacent neighbors. To maintain the optimal performance in the serial version of the simulator, we do not allow more sub-volumes than there are particles.

### S3. Reaction-Diffusion Pseudo code for the Serial Version

The following pseudo code describes the reaction-diffusion process in the Serial version of the simulator. Key constraints: Each molecule can participate in only one reaction per iteration step, even if it has multiple interfaces. It the molecule does not react, it moves as a rigid body. If a complex has multiple molecules, they are each allowed to try and react independently. If a complex does not react, it diffuses as a rigid body in that iteration step. Steric overlaps are prevented.

*Reaction-Diffusion Simulation (Serial Version)*

```
// Initialization

1. Generate all molecules and corresponding complexes

   Store molecules in vector: moleculeList

   Store complexes in vector: complexList
```

```
2. Calculate the cutoff_distance

   Divide simulation volume into sub_volumes based on cutoff_distance

   Store sub_volumes in vector: subVolList

   For each subVol in subVolList:

      Initialize vector<int> memberMolList

   For each molecule in moleculeList:

      Set molecule.mySubVolIndex to appropriate subVol index


// Main Simulation Loop

3. For iteration = 1 to max_interations:

   4. Update subVol memberships:

      For each subVol in subVolList:

         Clear subVol.memberMolList

      For each molecule in moleculeList:

         Update molecule.mySubVolIndex

         Append molecule.index to subVol.memberMolList

   5. Process zeroth-order and first-order reactions:

      Check and perform:

         - Molecule creation

         - Molecule destruction

         - Unimolecular state changes

         - Complex dissociation

   6. Calculate and store binding probabilities for all possible second-order bimolecular reactions.
```

*If the molecule interface already participated in a $0^{th}$ or $1^{st}$ order reaction, it can only have binding probabilities set to zero. This applies to all interfaces on the molecule. They must still be evaluated for second-order reactions, so that they will avoid overlap with reaction partners that are close by. The other elements of the complex (if the molecule is part of a complex) are not restricted from reacting, but are restricted from diffusing.

```
      For each subVol in subVolList:

         For each molecule1 in subVol.memberMolList:

            For each molecule2 in subVol.memberMolList:
```

```
            If binding_possible(molecule1, molecule2):
               Store binding information in molecule1 and molecule2
         For each subVol in adjacentSubVols:
            For each molecule2 in subVol.memberMolList:
               If binding_possible(molecule1, molecule2):
                  Store binding information in molecule1 and molecule2
   7. Perform second-order bimolecular reactions
      For each molecule in moleculeList:
         Compare the binding probabilities to a URN. If the probability is >URN, perform the reaction.
            *Note, that if this molecule already underwent a binding reaction, via another interface, then its probabilities for binding will all have been set to zero.
            Perform bimolecular reactions by associating molecule pair into their defined ‘bound’ orientation.
   8. Perform diffusion for unreacted complexes:
      For each molecule in moleculeList:
         If molecule has not undergone a 0th, 1st, or 2nd order reaction, or is not part of a complex that has undergone one of these reactions:
             Diffuse its complex as rigid body
             Ensure no steric overlaps occur with all molecules in its partner list, including molecules that have undergone reactions.
   9. Reset reaction information:
      For each molecule in moleculeList:
         Clear reaction status and information
   10. Update simulation time and collect data as needed

// End of main simulation loop
```

### S4. Description of the MPI Implementation of the Simulator

The MPI implementation of the simulator leverages distributed computing techniques to divide the simulation workload across multiple processors and nodes.

*1.1 Domain Decomposition*

a) The simulation volume is portioned along the x-axis, with each partition assigned to a distinct processor.
b) Each processor is responsible for a subset of the sub-volumes within its assigned region.
c) A processor corresponds to a single MPI rank.

*1.2 Processor Topology*

a) Processors are arranged in a linear topology along the x-axis.
b) For any given processor *n*:
    - If there is an adjacent processor on its left side, it is referred to as the "left neighbor processor", *n-1.*
    - Similarly, an adjacent processor on the right side is called the "right neighbor processor", *n+1.*

*1.3 Edge and Ghost Regions*

a) Edge Region: For a processor with a neighbor on one side, the sub-volumes at the boundary along the x-axis are designated as the "Edge region". The Edge region contains all the molecules that may interact with molecules in the neighboring processor's domain.
b) Ghost Region: A copy of a neighboring processor's Edge region. Thus, these sub-volumes are not owned by the processor. Ghost regions ensure the evaluation of reactions that occur across processor boundaries.

*1.4 Inter-Processor Communication*

The update of Ghost and Edge regions during the simulation is implemented using Message Passing Interface (MPI) functions.

*1.5 Extended Properties for Molecules and Complexes*

To facilitate efficient parallel processing and inter-processor communication, we introduce additional properties to the Molecule and Complex classes.

a) Global ID: Since each processor maintains its own local indexing system for molecules and complexes, global unique ID ensure unambiguous identification of entities throughout the distributed simulation environment.

b) Spatial Region Flags: Four Boolean properties are added to both Molecule and Complex on each processor:

i. isLeftGhost: True for a complex if any part of it is in the left Ghost region. True for a molecule in the left Ghost region. True for a molecule that is part of a complex in the left Ghost region, and that molecule is not in the left Edge region.
ii. isLeftEdge: True for a complex if any part of it is in the left Edge region. True for a molecule in the left Edge region. True for a molecule that is part of a complex in the left Edge region, and that molecule is not in the left Ghost region.
iii. isRightEdge: Same conditions as above.
iv. isRightGhost: Same conditions as above.

c) Region Assignment Consequences:

i. Molecules can only be 'True' for one of these four regions, or 'False' for all of them.
ii. Complexes can be 'True' for two regions if they span both an Edge and Ghost region.
iii. Every single molecule in a Complex that has a 'True' flag will either be assigned as a Ghost or Edge molecule. This includes molecules that extend out of both the ghost and edge sub-volumes. These molecules must be assigned a sub-volume on the physical processor, even though molecules beyond the ghost region exist outside of it. They are

assigned to the closest sub-volume by retaining the y and z index of the sub-volume, and setting the x index to 0.

d) Communication Tracking: A Boolean property receivedFromNeighbor is added to both Molecule and Complex classes. This property is used to track the loss of molecules and complexes from the observed processor during the inter-processor communication.

e) Communication Protocol:

a. Before communication: receivedFromNeighbor is set to false for all molecules and complexes in Edge and Ghost regions.

b. After receiving data from a neighbor processor: receivedFromNeighbor is set to true for all received entities.

c. Post-communication cleanup: Any molecule or complex in Edge or Ghost regions with receivedFromNeighbor == false is considered not received back from the neighbor and is deleted from the current processor.

### S5. Pseudo-code for Parallel reaction-diffusion simulator

```
1. Initialization:
  1.1 Initialize simulation domain and parameters
  1.2 Partition simulation volume along x-axis
  1.3 Assign partitions to available processors


2. Pre-Process Setup:
  For each processor:
   2.1 Initialize local sub-volumes, moleculeList, and complexList
   2.2 Assign global IDs to molecules and complexes
   2.3 Identify Edge regions based on the sub-volume indices. If the sub-volume x-index is 1, it is a left edge, if the sub-volume x-index is N-2, it is a right edge. The x-index is 0 for the left ghost-region. The x-index is N-1 for the right ghost-region.
   2.4 Set spatial region flags for molecule and complex, by looping over all molecules in the edge regions, and including all molecules that are shared by their parent complexes.


3. Main Simulation Loop:
  For each time step:
  3.2 Process Local Reactions:
      3.2.1 Check and perform zeroth-order and first-order reactions for all molecules in your physical processor, excluding the right edge region. Check
```

and perform these reactions for your left ghost region, and **exclude the right ghost region.**

3.2.2 Check for potential second-order bimolecular reactions for all particles in your physical processor. Include both the left edge and the right edge region. Include both the left ghost region and the right ghost region.

3.3 Divide Processor region:

Split your processor’s physical region into Left half and Right half from middle of x-axis. Create two lists, a left half list that contains the index of all molecules in the left half. A right half list contains the index of all molecules in the right half.

3.4 Process Left half:

3.4.1 Perform second-order bimolecular reactions for molecules in your physical left half and include your left ghost region. This includes all molecules that are assigned to the ghost region, even if it is due to being a member of a complex, and its physical position is outside of the ghost region. If a molecule associates, it has a new flag set to ‘isAssociated=true’.

3.4.2 Diffuse unreacted complexes (excluding left ghost region). If a molecule is part of a complex that is partially in the ghost region, it is not allowed to diffuse. Don’t diffuse complexes that spread across the left/right half and have not reacted. That will cause molecules to diffuse on the right half, even though they have not yet tried to react. Molecules can also diffuse into the right half, from the left half. Molecules can also diffuse into the left ghost region, from the left edge region.

3.4.3 Update sub-volume memberships and spatial region flags for all the molecules in your processor, including the ghost regions. The right ghost region should not have changed. This includes ghost molecules that are physically outside of your sub-volumes, but are part of a ghost-region complex.

3.5 Left-to-Right Communication:

If not last processor:

Set the receivedFromNeighbor to false for right edge and right ghost regions

If not first processor:

Send the left edge and ghost data to left neighbor processor

If not last processor:

Receive the data from right neighbor processor

Update the right ghost region and right edge region. That means that all molecules and complexes in these two regions either already had $0^{th}$, $1^{st}$, or $2^{nd}$ order reaction performed, or, they attempted but did not perform any reactions. Those reaction probabilities should all have been set to zero. Some of these molecules (most of those in the right ghost region) will already have diffused.

Delete unreceived molecules and complexes in right ghost/edge regions. This should only include molecules that were in your right ghost region, as they may have diffused beyond further to the right. It should not include any molecules in the right edge region, which could maximally have moved into the right ghost region, or if they diffused left you had assigned them to sub-volume 0 on the neighbor's processor.

3.6 Process Right half:

3.6.1 Perform second-order bimolecular reactions for all molecules in the right half based on the list 'RightHalfIndexes' established in 3.3. This excludes all molecules that diffused into the right half. This should exclude reactions to all molecules that were in the left half, as those reactions were already attempted.

This loop excludes all molecules in the right edge and ghost regions. However, molecules can attempt to react with any molecule in the right edge or right ghost region. These reactions are rejected if those molecules already underwent a 0th, 1st or 2nd order reaction. The isDissociated and isAssociated flags are used for this.

3.6.2 Diffuse unreacted complexes. All molecules that are in the right half, including molecules that are part of a complex that extends into the left half. All molecules that are in the right edge region that already diffused (via trajStatus::propagated = true flag) are not allowed to diffuse again. All molecules that are in the right ghost region and do not extend into the right edge region would have already diffused. Molecules that are in a complex that spans the edge and ghost might need to diffuse.

3.6.3 Update sub-volume memberships and spatial region flags

3.7 Right-to-Left Communication:

If not first processor:

Set the receivedFromNeighbor to false for left edge region and left ghost region

```
        If not last processor:
              Send the right edge and ghost data to right neighbor processor
        If not first processor:
              Receive the data from left neighbor processor
              Update the left ghost region and left edge region
              Delete unreceived molecules and complexes in left ghost/edge
        regions

    3.8 Post-Processing:
        3.8.1 Reset reaction information for all molecules
        3.8.2 Update sub-volume memberships, update spatial region flags for
  all molecules and complexes
        3.8.3 Update simulation time and collect data as needed

  4. Finalization:
        4.1 Merge results from all processors
        4.2   Generate final output
```

### S6. ODEs for Trimer Assembly Model

To model the trimer assembly system using ordinary differential equations (ODEs), we derive the rate of change in the concentrations of the species $[A]$, $[AA]$, $[AAA]$, and $[AAAc]$ based on the reaction rates and their stoichiometry. Here, $A$ represents the monomer, $AA$ is the dimer, $AAA$ is the open (non-closed) trimer, and $AAAc$ is the closed trimer. The reactions governing this system are as follows:

$$A + A \leftrightarrow AA, k1_{on}, k1_{off} \qquad (S1)$$

$$AA + A \leftrightarrow AAA, k2_{on}, k2_{off} \qquad (S2)$$

$$AAA \leftrightarrow AAAc, k_{close}, k_{open} \qquad (S3)$$

The ODEs for the above reactions:

$$\frac{d[A]}{dt} = -2k1_{on}[A]^2 + 2k1_{off}[AA] - 2k2_{on}[AA][A] + 2k2_{off}[AAA] \qquad (S4)$$

$$\frac{d[AA]}{dt} = k1_{on}[A]^2 - k1_{off}[AA] - 2k2_{on}[AA][A] + 2k2_{off}[AAA] \qquad (S5)$$

$$\frac{d[AAA]}{dt} = 2k2_{on}[AA][A] - 2k2_{off}[AAA] - k_{close}[AAA] + 3k_{open}[AAAc] \qquad (S6)$$

$$\frac{d[AAAc]}{dt} = k_{close}[AAA] - 3k_{open}[AAAc] \quad (S7)$$

Given the microscopic binding rate $k_a$ and unbinding rate $k_b$, the diffusion constant for monomer $D_A$, we have the equilibrium constant:

$$K_D = \frac{k_b}{k_a} = \frac{k1_{off}}{k1_{on}} = \frac{k2_{off}}{k2_{on}} \quad (S8)$$

We can get the macroscopic rates based on the microscopic rates:

$$k1_{on} = \left(\frac{1}{k_a} + \frac{1}{4\pi\sigma(2D_A)}\right)^{-1} \quad (S9)$$

$$k2_{on} = \left(\frac{1}{k_a} + \frac{1}{4\pi\sigma(D_A + D_{AA})}\right)^{-1} \quad (S10)$$

$D_{AA} = \frac{D_A}{2}$. The loop closure rate is a first-order reaction rate given by

$$k_{close} = k_a C_0 exp\left(-\Delta G_{coop}/k_B T\right) \quad (S11)$$

$$k_{open} = k_b \quad (S12)$$

Where $C_0 = 1M$ is the standard state concentration, and $\Delta G_{coop}$ can introduce positive or negative cooperativity into the loop closure reaction. We use $\Delta G_{coop} = 0$ for no cooperativity.

### SUPPLEMENTAL FIGURES

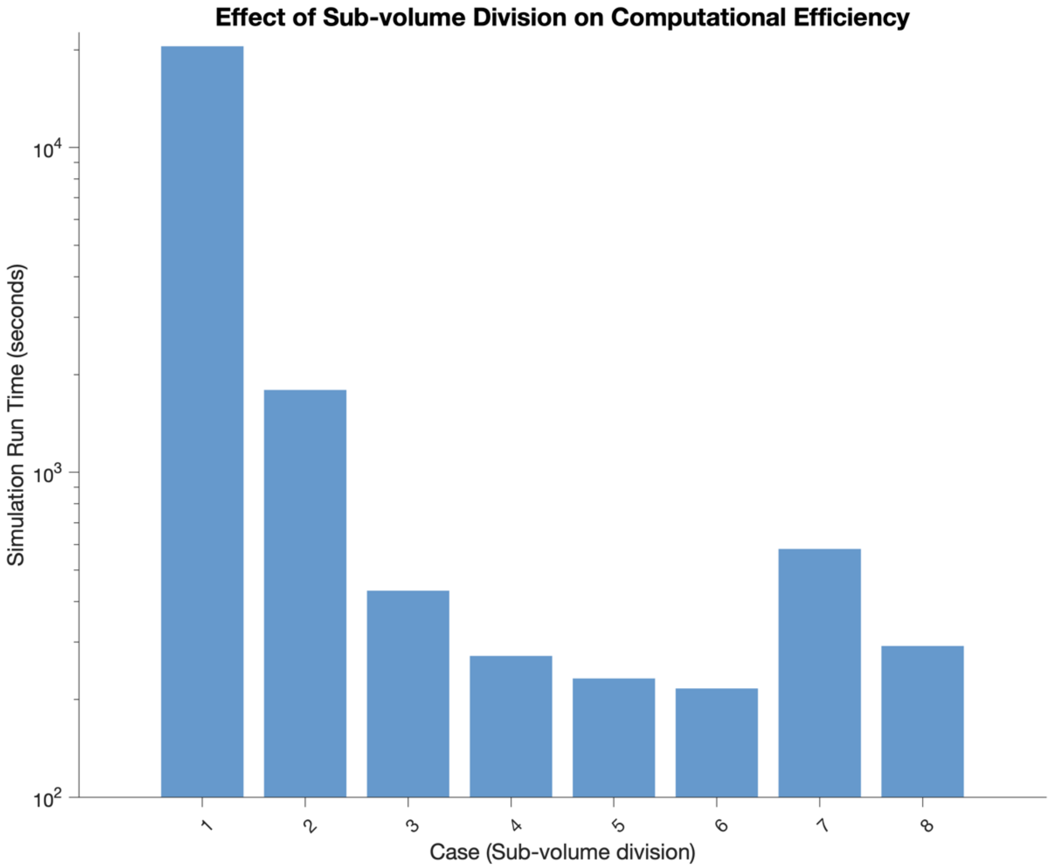


**Fig S1. Effect of sub-volume division on computational efficiency in 3D bimolecular reaction simulations.** This figure illustrates the effect of different sub-volume division strategies on the computational efficiency of simulating the bimolecular reaction A + B <-> A.B in 3D. Simulations were conducted in a system of size [642.94, 642.94, 642.94] nm, with initial copy numbers of 8000 for both A and B molecules. Reaction parameters were set to ka = 20 $nm^3/\mu s$, kb = 10000/s, $\sigma$ = 2nm, and D = 10 $nm^2/\mu s$, with a time step of 0.01 $\mu$s for 10000 iterations. Eight cases with varying sub-volume divisions were tested: Case 1 [4,4,4], Case 2 [10,10,10], Case 3 [20,20,20], Case 4 [30,30,30], Case 5 [40,40,40], Case 6 [50,50,50], Case 7 [100,100,100], and Case 8 [121, 14, 14]. All simulations were performed using a single processor. One simulation is run for each case. Case 8 is designed to demonstrate the impact of asymmetric division. The results reveal that increasing sub-volume division initially improves computational efficiency, with an optimal point, beyond which efficiency decreases. Asymmetric division has little effect on the computational efficiency.

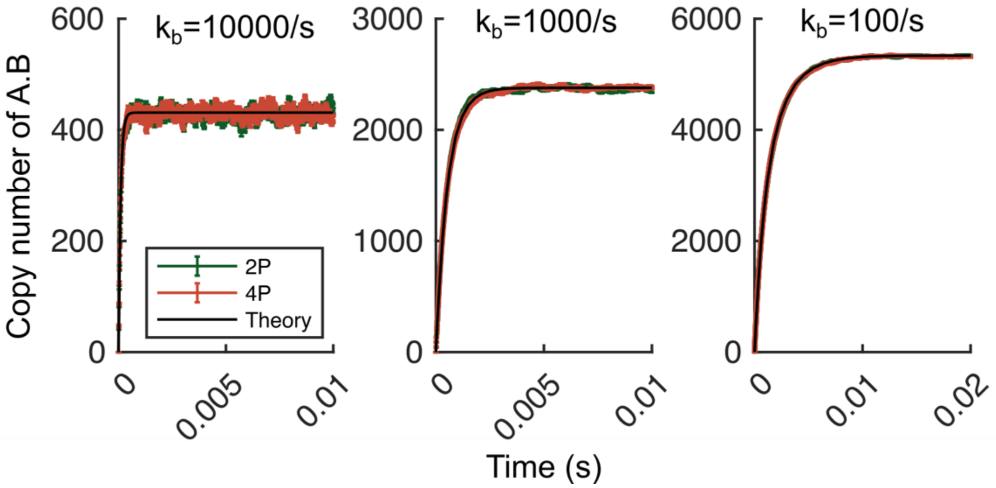


**Fig S2. Validation of accuracy for the parallel implementation at varying binding constants (3D bimolecular reaction simulations).** This figure compares the accuracy of the parallel implementation for simulating the bimolecular reaction A + B <-> A.B in 3D with the ODE solutions. Simulations were conducted in a system of size [642.94, 642.94, 642.94] nm, with initial copy numbers of 8000 for both A and B molecules. $\sigma$ = 2nm. ka = 20 $nm^3/\mu s$. D = 10 $nm^2/\mu s$ for A and B. time step = 0.1 $\mu s$. Case 1: kb = 10000/s; Case 2: kb = 1000/s; Case 3: kb = 100/s. Each case was run using 2 processors and 4 processors, and the results averaged across 6 traces. Simulation results are presented as the mean ± standard error (SE). Both the kinetic and equilibrium match the theoretical expectations.

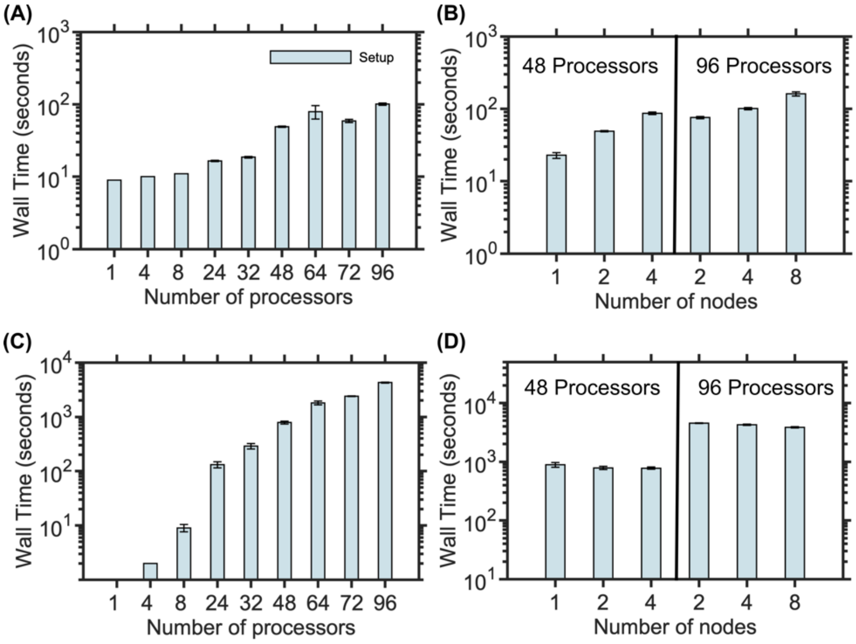


**Fig S3. Setup time for 3D bimolecular reaction simulations in Fig 3.** (A, B) strong scaling benchmark system. (C, D) weak scaling benchmark system.

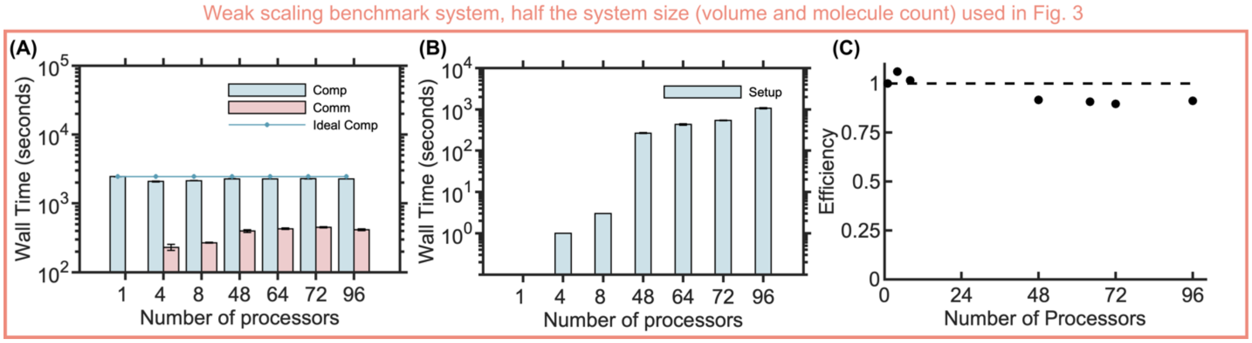


**Fig S4. Weak scaling benchmark with half size compared to main text Fig 3.** (A) Computational and communication times. (B) Setup time. (C) Efficiency.
